# Single cell analysis of lincRNA expression during human blastocyst differentiation identifies ***TERT* ^*(+)*^** multi-lineage precursor cells

**DOI:** 10.1101/084087

**Authors:** Jens Durruthy-Durruthy, Mark Wossidlo, Vittorio Sebastiano, Gennadi Glinsky

## Abstract

Chromosome instability and aneuploidies occur very frequently in human embryos, impairing proper embryogenesis and leading to cell cycle arrest, loss of cell viability, and developmental failures in 50-80% of cleavage-stage embryos. This high frequency of cellular extinction events represents a significant experimental obstacle challenging analyses of individual cells isolated from human preimplantation embryos. Here, we carried out single cell expression profiling analyses of 241 individual cells recovered from 32 human embryos during the early and late stages of viable human blastocyst differentiation. Classification of embryonic cells was performed solely based on expression patterns of human pluripotency-associated transcripts (*HPAT*), which represent a family of transposable element-derived lincRNAs highly expressed in human embryonic stem cells (hESCs) and regulating nuclear reprogramming and pluripotency induction. We then validated our findings by analyzing 1,708 individual embryonic cells recovered from more than 100 human embryos and 259 mouse embryonic cells at different stages of preimplantation embryogenesis. Our experiments demonstrate that segregation of human blastocyst cells into distinct sub-populations based on single-cell expression profiling of just three *HPATs* (*HPAT-21; -2;* and *-15*) appears to inform key molecular and cellular events of naïve pluripotency induction and accurately captures a full spectrum of cellular diversity during human blastocyst differentiation. *HPAT*’s expression-guided spatiotemporal reconstruction of human embryonic development inferred from single-cell expression analysis of viable blastocyst differentiation enabled identification of *TERT*^(+)^ sub-populations, which are significantly enriched for cells expressing key naïve pluripotency regulatory genes and genetic markers of all three major lineages created during human blastocyst differentiation. Results of our analyses suggest that during early stages of preimplantation embryogenesis putative immortal multi-lineage precursor cells (iMPCs) are created, which then differentiate into trophectoderm, primitive endoderm and pluripotent epiblast lineages. We propose that cellular extinction events in cleavage-stage embryos are triggered by premature activation of HPAT lincRNAs reflecting failed iMPC’s creation attempts.

**Highlights:**

- Single cell analysis of 1,949 human & 259 mouse embryonic cells
- Identification of 5 most abundant HPAT lincRNAs in viable human blastocysts
- Expression profiling of just 3 lincRNAs captures cellular diversity of human blastocysts
- Identification & characterization of *TERT*^*(+)*^ multi-lineage precursor cells
- *MTTH/HPAT* lincRNAs regulatory axis of naïve pluripotency induction *in vivo*

## Introduction

Precise spatio-temporal activation of species-specific transposable elements (TrE), and particularly of regulatory sequences derived from human endogenous retroviruses (HERVs), have been associated with the induction and maintenance of human pluripotency, functional identity and integrity of naïve-state embryonic stem cells, and anti-viral resistance of the early-stage human embryos [1–17]. These studies identified long intergenic non-coding RNAs (lincRNAs) derived from HERVs as the reliable genetic markers and critically important regulators of the pluripotent state in human cells [8; 14; 15; 17]. Active expression of lincRNAs encoded by specific HERVs is required for pluripotency maintenance in hESC [8; 17] and plays essential functional roles in naïve-like hESC [15; 16], development of human preimplantation embryos [16; 17], and pluripotency induction during nuclear reprogramming [9; 10; 15; 17]. Individual members of one functionally and structurally-related family of TrE-derived lincRNAs, which are highly expressed in hESC and termed human pluripotency-associated transcripts (*HPATs*), have been shown to affect the regulation of nuclear reprogramming and pluripotency induction during preimplantation embryogenesis [17]. Despite this significant progress facilitated by single-cell analyses of hESCs and human embryos, our understanding of how the earliest lineage commitments are regulated and individual cell fate decisions are executed remains limited. In particular, the expression dynamics and biological roles of TrE-derived lincRNAs in these processes are not fully understood.

Several methodological hurdles and fundamental problems complicate the analysis of human preimplantation embryogenesis. Chromosome instability is common in the early-stage human embryonic development and aneuploidies are observed in 50-80% of cleavage-stage human embryos leading to cell cycle arrest, impaired viability of embryonic cells’ and developmental failures [19–23]. Therefore, without the application of reliable prospective controls of the viability of human embryonic cells, the likelihood of studying a significant fraction of dying human embryonic cells is high and should approach the 50 - 80% probability. A systematic application of non-invasive assays that accurately predict the likelihood of viable blastocyst development during evaluation of the mature oocytes and/or zygotes should help to address this limitation. It has been shown that the analysis of zygote viscoelastic properties as well as cytokinesis timing in the early stages of human embryo development can predict viable blastocyst formation with >90% precision, 95% specificity and 75% sensitivity [23; 46]. Another potentially significant limitation of single cell studies of human embryos is the use of the timeline after fertilization as end-point for isolating individual human embryonic cells. This approach would likely generate developmentally heterogeneous mixes of single cells isolated from human embryos that are not fully synchronized and will have varying proportions of dying versus viable cells at any given developmental time point.

To address these limitations, we implemented an experimental approach ensuring the high likelihood of studying truly viable human embryonic cells and employing the strict morphological criteria for segregation of viable blastocyst cells into the early and late blastocyst developmental stages defined by classic embryological criteria [18]. Using this approach, single-cell gene expression profiling was successfully carried-out in 241 individual cells recovered from early and late human blastocysts to delineate the spatiotemporal dynamics of gene expression changes during human blastocyst differentiation [18]. This experimental system faithfully distinguished all three lineages that form the human blastocyst and facilitated development of a three-dimensional *in silico* model of the inner cell mass and trophectoderm, in which individual cells were mapped into distinct expression domains corresponding to the lineage differentiation in human blastocysts [18]. This *in situ* model was then used to design the first web-based online tool to study early cell fate decisions in the human blastocyst, which is available online at http://web.stanford.edu/~sunilpai/HumanBlastocystViewer.html (Firefox/Chrome compatible).

Here, we utilized this experimental system to perform analysis of TrE-derived lincRNAs within the context of the high-complexity single-cell expression profiling matrix comprising more than 20,000 features and capturing the expression of 89 genes in 241 individual cells recovered from early and late differentiation stages of viable human blastocysts. We validated our experimental findings and novel mechanistic models by *in silico* gene expression profiling analyses of four independent validation data sets comprising of 1,708 individual embryonic cells recovered from more than 100 human embryos at different stages of preimplantation embryogenesis [12; 13; 34; 44}. The major novel observations reported in this contribution are:

i) Our experiments identify five most abundant *HPATs* (*HPAT21/15/5/2;3*) which were detected in 25% to 97% of viable human blastocyst cells;
ii) We demonstrate that segregation of human blastocyst cells into distinct sub-populations based on a single-cell expression profiling of just three *HPATs* (*HPAT21/2/15*) appears to inform key molecular and cellular events of naïve pluripotency induction and accurately captures the entire spectrum of cellular diversity during human blastocyst differentiation;
iii) We observe that during human embryogenesis, telomerase-positive multi-lineage precursor cells are created to then differentiate into trophectoderm, primitive endoderm and pluripotent epiblast lineages.

## Results

### HPAT lincRNA expression-guided classification of human blastocyst cells

Using a hybrid RNA sequencing technique we have previously identified and characterized 23 novel TrE-derived lincRNAs that are highly expressed in hESC and termed human pluripotency-associated transcripts (*HPATs*) [17; 36]. We have developed and implemented qPCR-based assays for sixteen HPATs [17; 18] and thought to begin our analytical effort by focusing on the exploration of the expression patterns of these sixteen TrE-derived lincRNAs during human blastocyst differentiation (Fig. 1). This analysis identifies five TrE-derived lincRNAs (*HPAT21; HPAT15; HPAT5; HPAT2;* and *HPAT3*) which were observed most frequently and detected in at least 25% of analyzed human blastocyst cells (Supplemental Fig. S1). Notably, three out of these five lincRNAs were recently implicated in regulation of nuclear reprogramming and pluripotency induction during human preimplantation embryo development [17]. Expression of *HPAT21,* the most abundant lincRNA, was detected in 92% of cells recovered from the early blastocysts and 100% of the late blastocyst cells, whereas expression of the second-ranked lincRNA *HPAT15* was observed in 56% and 48% of the early and late blastocyst cells, respectively. Notably, the expression patterns of the vast majority (94%) of lincRNAs did not manifest significant changes in the early and late blastocyst cells (Supplemental Fig. S1), indicating that xpression of individual TrE-derived lincRNAs remains relatively stable and may serve as a reliable marker of distinct sub-populations emerging during human blastocyst differentiation. The detailed analysis of expression patterns of all individual lincRNAs in 241 individual human blastocyst cells revealed that there is only one cell manifesting *HPAT21/HPAT2/HPAT15*-null phenotype and remaining human blastocyst cells express varying combinations of these three lincRNAs (Fig. 1B). The results of the present analyses suggest that expression profiles of just three lincRNAs capture a full spectrum of cellular phenotypes created during differentiation of human blastocysts. It was of interest to determine whether detailed analysis of sub-populations of cells defined by the expression patterns of these 3 genetic markers would inform events of naïve pluripotency induction and faithfully describe the evolution of cellular diversity during human blastocyst differentiation.

**Fig. 1.**
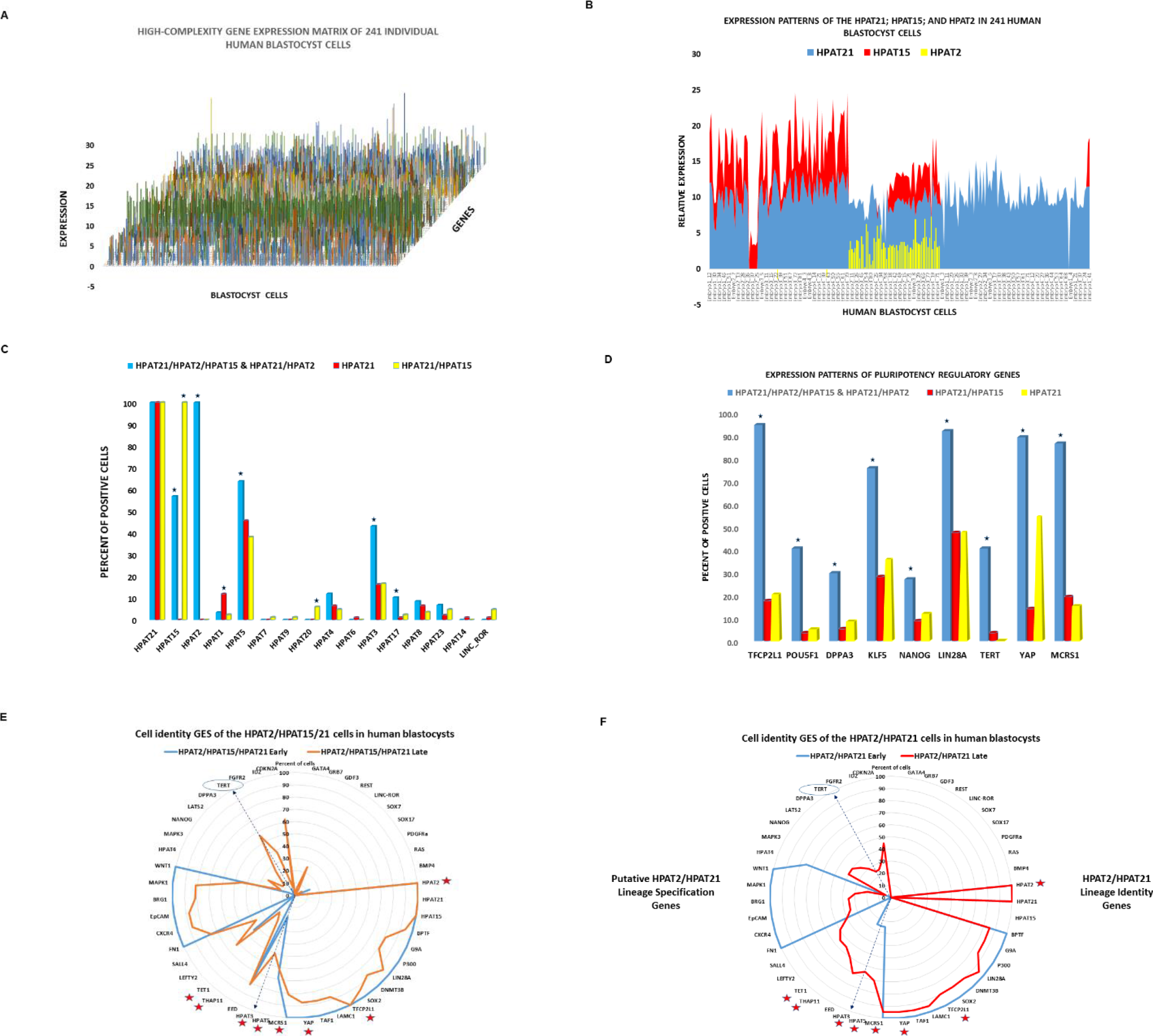
Single-cell gene expression analysis of HPAT lincRNAs captures the entire spectrum of a cellular diversity during human blastocyst differentiation. **(A)** A high complexity expression matrix of 89 genes expression of which were measured in 241 individual human blastocyst cells. **(B)** Expression profiles of the *HPAT21; HPAT15;* and *HAPT2* lincRNAs define major sub-populations of human blastocyst cells. A color-coded stacked area plot depicts relative expression values of designated HPAT lincRNAs in each individual blastocyst cell. Note that there is only one cell manifesting *HPAT21/HPAT2/HPAT15*-null phenotype. **(C)** Expression patterns of HPAT lincRNAs in distinct sub-populations of human blastocyst cells. Stars designate sub-populations harboring the significantly enriched numbers of cells expressing defined genetic markers. P values were estimated using two-tailed Fisher’s exact test and reported in the Supplemental Table S2. **(D)** Expression patterns of pluripotency regulatory genes in distinct sub-populations of human blastocyst cells. Stars designate sub-populations harboring the significantly enriched numbers of cells expressing defined genetic markers. **(E)** Cell identity gene expression signature (GES) of the *HPAT21* ^*(+)*^ *HPAT15* ^*(+)*^ *HPAT2* ^*(+)*^ cells recovered from human blastocysts. Stars designate transcripts encoded by the genomic loci that were implicated in the regulation of the human pluripotency state. Positions of colored lines’ heights inside the circle reflect the percentage of cells within a population that express transcripts encoded by the corresponding genes names of which are listed on the circle. **(F)** Cell identity gene expression signature (GES) of the *HPAT21* ^*(+)*^ *HPAT15* ^*(−)*^ *HPAT2* ^*(+)*^ cells recovered from human blastocysts. Stars designate transcripts encoded by the genomic loci that were implicated in the regulation of the human pluripotency state. Positions of colored lines’ heights inside the circle depict the percentage of cells within a population that express transcripts encoded by the corresponding genes listed on the circle.

### Single-cell expression profiling of HPAT expression-defined sub-populations during human blastocyst differentiation

Early and late blastocyst cells were segregated into distinct sub-populations based on expression patterns of three lincRNAs (*HPAT21; HPAT2;* and *HPAT15*) and gene expression signatures (GES) of these *HPAT* expression-defined sub-populations of human blastocyst cells were identified and analyzed. The results of these analyses are shown in the Figs 1–5. All human blastocyst cells are segregated into sub-groups based on common *HPAT*s’ expression patterns (Figs. 1 and 2E). The numbers of cells comprising the corresponding *HPAT*’s expression-defined sub-populations of early (Fig. 2F) and late (Fig. 2G) blastocysts are shown.

**Figure 2.**
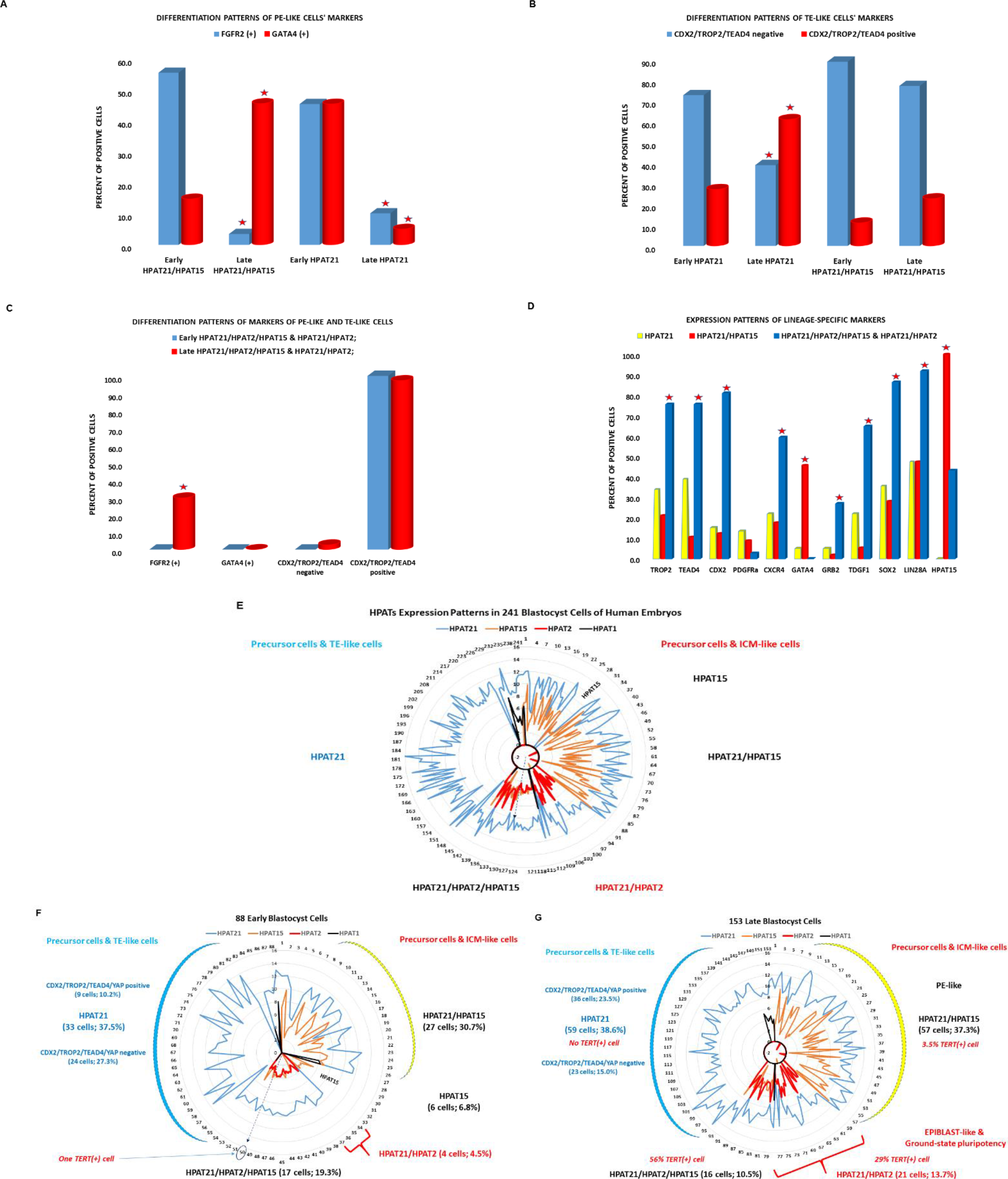
Single-cell analysis of expression patterns of the lineage-specific genetic markers in HPAT lincRNA expression-defined sub-populations during human blastocyst differentiation. Expression changes of designated lineage-specific genetic markers (see text for details) were evaluated in individual cells comprising the corresponding HPAT expression-defined sub-populations that were recovered from the early-stage and late-stage human blastocysts. Differentiation patterns in human blastocyst cells of genetic markers of PE-like cells are shown in (A) and (C). Differentiation patterns of genetic markers of TE-like cells are shown in (B). Note that HPAT21 ^(±)^ HPAT15 (± HPAT2 ^(-)^ cells manifest gene expression differentiation patterns typical of PE-like cells (A), while HPAT21 ^(±)^ HPAT15 ^(-)^ HPAT2 ^(-)^ cells exhibit gene expression differentiation patterns typical of TE-like cells (B). In contrast, no gene expression differentiation patterns resembling either PE-like or TE-like cells were observed in HPAT2 ^(±)^ cell sub-populations (C). D) HPAT2 ^(±)^ cell sub-populations manifest significant enrichment of cells expressing genetic markers of all three major lineages created during human blastocyst differentiation. (E) HPAT lincRNA expression patterns in 241 individual human blastocyst cells. Each number on the circle designates one randomly placed blastocyst cell. (F) HPAT lincRNA expression patterns in 88 individual cells recovered from the early-stage human blastocysts. Each number on the circle designates one randomly placed early-stage blastocyst cell. (G) HPAT lincRNA expression patterns in 153 individual cells recovered from the late-stage human blastocysts. Each number on the circle designates one randomly placed late-stage blastocyst cell. In the Figs. 2E-G each number on the circle corresponds to a single blastocyst cell. The colored lines inside the circle depict the corresponding *HPAT* lincRNAs and positions of the lines’ heights reflect the *HPAT* lincRNAs’ expression values in corresponding cells. All human blastocyst cells are segregated into sub-groups based on common *HPATs’* expression patterns (Fig. 1). The numbers of cells comprising the corresponding *HPAT’s* expression-defined sub-populations of the early (Fig. 2F) and late (Fig. 2G) blastocysts are shown. The statistical significance of the observed expression changes between sub-populations or within a sub-population during differentiation were estimated based on comparisons of the numbers of positive and negative cells using two-tailed Fisher’s exact test. Stars designate sub-populations harboring the significantly different numbers of cells expressing defined genetic markers. P values reported in the Supplemental Table S2.

The overall structure of the *HPAT*’s expression-defined sub-populations remains similar during the transition from the early to late blastocysts (Figs. 1; 2E-2G). The only sub-population that appears to exist exclusively among the early blastocyst cells is represented by the single-positive *HPAT21* ^*(−)*^ *HPAT2* ^*(−)*^ *HPAT15* ^*(+)*^ cells (Figs. 1B, 2E-2G) comprising a minority of the early blastocyst cell population (in total, 6 cells; 6.8%). Another single-positive sub-population is represented by the ninety-two *HPAT21* ^*(+)*^ *HPAT2* ^*(−)*^ *HPAT15* ^*(−)*^ cells comprising 37.5% (33 cells) and 38.6% (59 cells) of the early and late blastocyst cell populations, respectively (Figs. 2E-2G). The remaining sub-populations of the both early and late blastocysts are represented by the triple-positive *HPAT21* ^*(+)*^ *HPAT2* ^*(+)*^ *HPAT15* ^*(+)*^ cells and two distinct double-positive sub-populations of *HPAT21* ^*(+)*^ *HPAT2* ^*(+)*^ *HPAT15* ^*(−)*^ and *HPAT21* ^*(+)*^ *HPAT2* ^*(−)*^ *HPAT15* ^*(+)*^ cells (Figs. 1; 2E–2G). Notably, expression patterns of seven TrE-derived lincRNAs appear significantly different in these sub-populations (Fig. 1C) and reliably distinguish three sub-populations: *HPAT21* ^*(+)*^ *HPAT2* ^*(+)*^ *HPAT15* ^*(+)*^*& HPAT21* ^*(+)*^ *HPAT2* ^*(+)*^ *HPAT15* ^*(−)*^ cells; *HPAT21* ^*(+)*^*HPAT2* ^*(−)*^ *HPAT15* ^*(−)*^ cells; and *HPAT21* ^*(+)*^ *HPAT2* ^*(−)*^ *HPAT15* ^*(+)*^ cells (Fig. 1). Significantly, the expression of the pluripotency regulatory genes (also defined as the genetic markers of the epiblast-like sub-populations in human blastocysts) appears markedly enriched in the *HPAT21* ^*(+)*^ *HPAT2* ^*(+)*^ *HPAT15* ^*(+)*^ *& HPAT21* ^*(+)*^ *HPAT2* ^*(+)*^ *HPAT15* ^*(−)*^ cells compared with the *HPAT21* ^*(+)*^ *HPAT2* ^*(−)*^ *HPAT15* ^*(−)*^ and *HPAT21* ^*(+)*^ *HPAT2* ^*(−)*^ *HPAT15* ^*(+)*^ cells (Fig. 1D). We focused our subsequent analytical effort on detailed characterization of the following three HPAT expression-defined sub-populations of human blastocyst cells:

i) *HPAT21* ^*(+)*^ *HPAT2* ^*(−)*^ *HPAT15* ^*(−)*^ cells, which were designated for clarity as the single-positive *HPAT21* cells (*spHPAT21*);
ii) *HPAT21* ^*(+)*^ *HPAT2* ^*(−)*^ *HPAT15* ^*(+)*^ cells, which were designated for clarity as the double-positive *HPAT15* cells (*dpHPAT15*);
iii) *HPAT21* ^*(+)*^ *HPAT2* ^*(+)*^ *HPAT15* ^*(−)*^ & *HPAT21* ^*(+)*^ *HPAT2* ^*(+)*^ *HPAT15* ^*(+)*^ cells, which were designated for clarity as the *HPAT2* positive cells (*HPAT2pos*);

Single-cell expression profiling experiments provide the unique opportunity to characterize the distinct sub-populations of cells based on the relative prevalence within a sub-population of individual cells expressing the particular sets of genetic markers. We thought to utilize this approach, rather than comparisons of mean expression values, reasoning that applications of these single cell analysis-enabled metrics would be more informative in distinguishing different sub-populations by performing the formal statistical evaluation of differences in the relative prevalence of cells expressing the particular sets of genetic markers within distinct *HPAT* expression-defined sub-populations of human blastocysts. The *HPAT2pos* sub-populations manifest similar gene expression signatures that remain relatively stable during the blastocyst differentiation (Figs. 1E & F). This is particularly evident for *HPAT21* ^*(+)*^ *HPAT2* ^*(+)*^ *HPAT15* ^*(+)*^ cells (Fig. 1E) indicating that these cells do not experience dramatic large-scale changes in gene expression during the transition from the early to late blastocyst stages. Most visible common changes observed in these two sub-populations during blastocyst differentiation were increase in the number of *TERT* ^*(+)*^ and *HPAT3* ^*(+)*^ cells (marked by the arrows in the Figs. 1E & 1F). Overall, the functional identities of genes comprising the gene expression signatures of the *HPAT2pos* sub-populations appear highly consistent with the putative biological role of these cells as the ground-state pluripotency precursors defined as the epiblast-like cells (Figs. 1D-1F).

We observed that structures of the *HPAT21* ^*(+)*^ *HPAT2* ^*(+)*^ *HPAT15* ^*(−)*^ sub-populations manifest changes of cellular compositions during the transition from early to late blastocyst stages (Fig. 1F). These changes appear to reflect the diversification of cellular compositions within early- and late-stage sub-populations, which is characterized by the relative losses of cells expressing the *FN1; CXCR4; EpCAM; BRG1; MAPK1; WNT1; HPAT4* lincRNA transcripts and the relative gains of cells expressing the *HPAT5; HPAT3; EED; THAP11; TET1; LEFTY2; SALL4; MAPK3; NANOG; LATS2; DPPA3; TERT; FGFR2; ID2; CDKN2A* transcripts (Fig. 1F). Notably, similar although apparently less profound changes were observed within the early- and late-stage sub-populations of the *HPAT21* ^*(+)*^ *HPAT2* ^*(+)*^ *HPAT15* ^*(+)*^ triple-positive cells (Fig. 1E).

### Analysis of expression profiles of genetic markers of distinct lineages created during human blastocyst differentiation

In contrast to the *HPAT2pos* cells, we observed clear evidence that other *HPAT*’s expression-defined sub-populations appear to display the gene expression changes which were previously identified as genetic elements of distinct differentiation programs during the human blastocyst development (Fig. 2). The validity of the proposed lineage assignments to the *HPAT* expression-defined sub-populations in human blastocyst were assessed using single-cell expression profiling data of genes specifically expressed in the trophectoderm (*TROP2, CDX2, TEAD4*); expression profiling of known markers of the pluripotent epiblast (*LIN28A* and *TDGF1*) and known markers of primitive endoderm (*GRB2* and *GATA4*) [24–26] as well as additional sets of the human blastocyst lineage-specific genetic markers identified in the recent study [18]. It has been documented that primitive endoderm (PE) cells express high levels of the *FGFR2* and low levels of the *GATA4* at the early-stage blastocyst differentiation [18]. During the transition from the early to late blastocyst stages, the *FGFR2* expression in the PE cells is markedly diminished, while expression of *GATA4* is concomitantly significantly increased [18]. These observations are highly consistent with previous reports demonstrating that FGFR2 is expressed by PE progenitor cells and is an essential signaling factor for PE differentiation in the mouse [24; 26; 27]. FGFR2-expressing PE progenitor cells bind to FGF4, which is released by epiblast cells, resulting in repression of *NANOG* and activation of *GRB2* and *MAPK* genes in PE progenitor cells, which ultimately leads to the induction of the mature PE specific marker GATA4 [28; 29]. Remarkably, the *dpHPAT15* ^*(+)*^ cells, in striking contrast to other *HPAT* expression-defined sub-populations recovered from early and late blastocysts, recapitulated exactly the PE-like patterns of *FGFR2* and *GATA4* expression changes during human blastocyst differentiation (Figs. 2A-2C). These data suggest that the *dpHPAT15* cells resemble the PE-like sub-population of human blastocysts. This conclusion is in a good agreement with the recent observation that *HPAT15* lincRNA appeared to be predominantly expressed in mature PE cells [18].

Expression analysis of markers of trophectoderm (TE) cells demonstrates that the *spHPAT21* cells recovered from the early-stage blastocysts contain only 27% of *CDX2/TROP2/TEAD4*-positive cells (Fig. 2B). In contrast, the *CDX2/TROP2/TEAD4*-positive cells became a dominant sub-population (67%) among the *spHPAT21* cells recovered from late-stage blastocysts. This marked enrichment for cells expressing TE-like genetic markers during blastocyst differentiation appears specific to the *spHPAT21* sub-population since it has not been observed in other *HPAT* expression-defined sub-populations recovered from early- and late-stage human blastocysts (Figs. 2B & 2C). Collectively, these data indicate that the *spHPAT21* cells resemble the TE-like lineage of human blastocysts.

Previous analyses demonstrate that cells expressing genetic markers of the pluripotent epiblasts are significantly enriched within the *HPAT2pos* sub-populations (Fig. 1). Strikingly, analysis of expression patterns of lineage-specific markers in *HPAT* expression-defined sub-populations recovered from late-stage human blastocysts revealed that cells expressing multiple markers of both TE and PE lineages appear significantly more prevalent among the *HPAT2pos* sub-populations (Fig. 2D). These results demonstrate that *HPAT2pos* cells express markers of all three major lineages created during blastocyst differentiation (Fig. 2D).

Collectively, our observations strongly argue that during human blastocyst differentiation, the unique sub-population of *HPAT2pos* human embryonic cells is created that could be defined as putative multi-lineage precursor cells. It is tempting to speculate that subsequent specialization and differentiation of *HPAT2pos* cells fuels the emergence of the trophectoderm, primitive endoderm and epiblast lineages.

### Expression patterns of human naïve pluripotency regulatory genes during human blastocyst differentiation

Several recent studies reported identification of genes that play essential role in induction, reset, and/or maintenance of the naïve pluripotent state in human cells [18; 30–32]. Forced expression of these genes induce or reset the naïve pluripotent state in human cells whereas their targeted silencing markedly affect the naïve pluripotency phenotypes. We observed that blastocyst cells expressing the genes encoding all these naïve pluripotency inducers are markedly enriched within the *HPAT2pos* sub-populations (Fig. 3A-B), particularly among the cells recovered from late-stage blastocysts.

To further test these observations, we identified the consensus gene expression signature (GES) of the induced naïve pluripotent state based on microarray analyses reported in recent studies, which utilized unique and markedly different protocols of naïve pluripotency induction, reset, and maintenance [30; 31]. This consensus GES of the human induced naïve pluripotent state comprises 13 genes, expression of which was consistently significantly altered in naïve pluripotent cells compared with the primed pluripotent cell cultures resulting in highly correlated profiles of their gene expression changes (Fig. 3C). We thought to utilize this consensus GES to further test the hypothesis that *HPAT2pos* sub-populations of human blastocyst cells may contain putative immediate precursors of naïve pluripotent cells. Strikingly, we observed that human blastocyst cells expressing 11 of 13 (85%) of these genes are markedly enriched within the *HPAT2pos* sub-populations compared with other cells recovered from human blastocysts (Fig. 3D).

**Figure 3.**
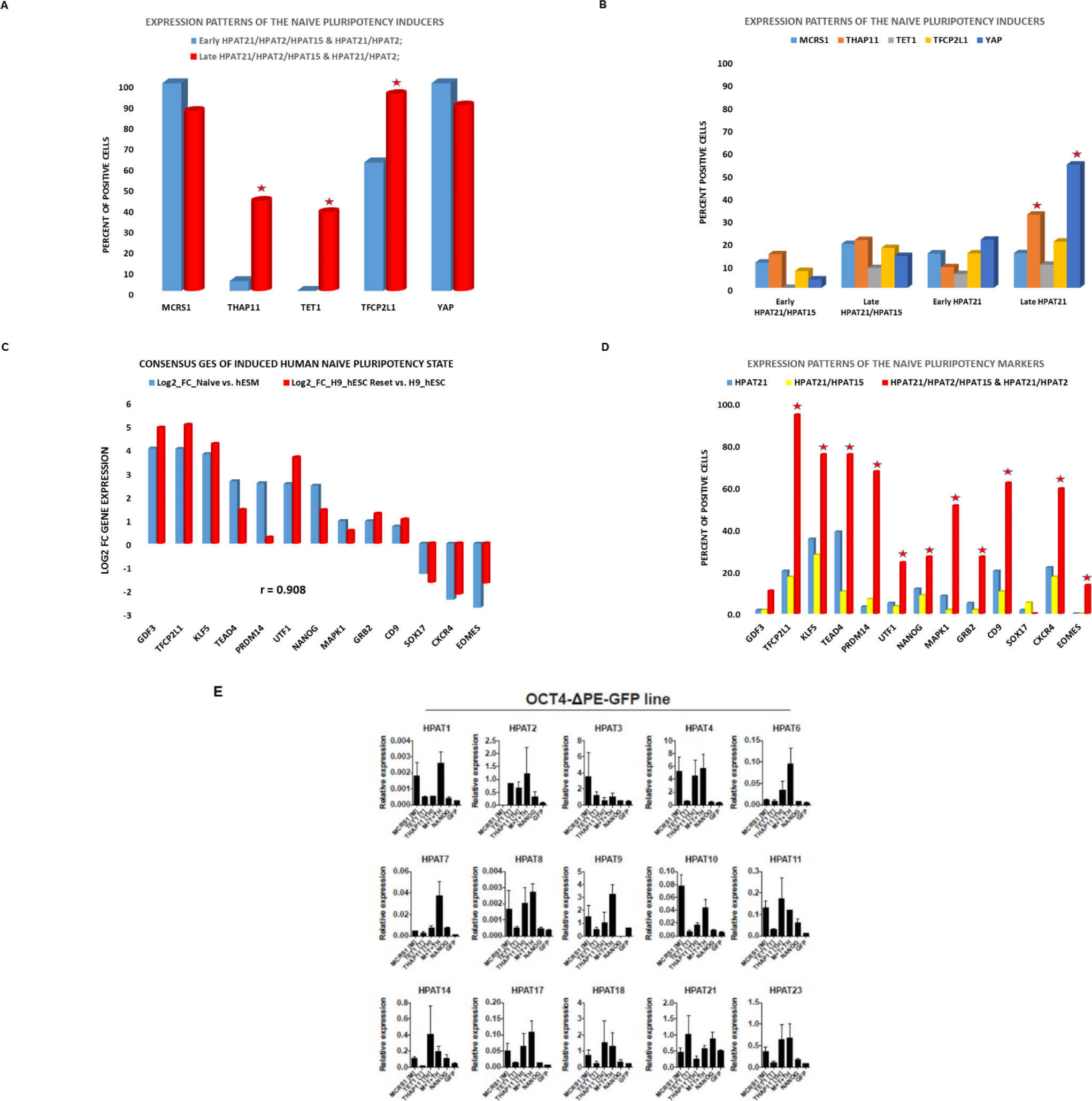
Single-cell next generation sequencing-defined dynamics of expression changes of LTR7/HERVH loci of the HPAT lincRNA genes during human preimplantation embryonic development. Panels (A-E) show the patterns of expression changes defined by the mean expression values of transcripts encoded by the designated loci at corresponding stages of human embryogenesis. Panels (F-I) report the expression values of designated transcripts in 156 individual cells recovered from the human embryos at the designated developmental stages. Results of the additional analyses are reported in the Supplemental Fig. S2.

### Activation of the *MTTH/HPAT* lincRNA regulatory axis during human embryogenesis

Previous reports demonstrated that expression of HERVH-derived lincRNAs [8; 15] and HPAT lincRNAs [17] in human pluripotent cells is regulated by a combination of transcription factors defined as a core pluripotency regulators (Fig. 6I). We recently showed [18] that expression of the three protein-coding genes *MCRS1, TET1, THAP11 (MTTH),* and *NANOG,* was concomitantly highly upregulated in the early precursor epiblast cells recovered from human blastocysts and demonstrated that forced expression of these three genes (*MTTH*) induces the naïve pluripotency-like phenotype in human cells. Notably, cells expressing all three *MTTH* genes implicated in the naïve pluripotency induction in human cells are significantly enriched among the *HPAT2pos* sub-populations recovered from late-stage human blastocysts (Fig. 3A). Based on these observations, we thought to utilize the *MTTH* model of induced human naïve pluripotent cells to determine whether the expression of HPAT lincRNAs is affected in human cells that acquired the naïve pluripotency phenotype. Strikingly, we observed (Fig. 3E) that the expression of a majority of HPAT lincRNAs was significantly increased in *MTTH*-induced human naïve pluripotent cells which were engineered in accordance with the previously reported protocol [18]. Of note, expression of some HPAT lincRNAs appears significantly up-regulated in *MCRS1*-overexpressing cells (Fig. 3E), suggesting that the *MCRS1* gene product may regulate the expression of HPAT lincRNAs perhaps by relieving the DAXX protein-mediated transcriptional repression via direct protein-protein interaction and sequestering DAXX to the nucleolus [33]. These findings are consistent with the hypothesis that the *MTTH*-induced naïve pluripotent state is associated with marked up-regulation of the HPAT lincRNAs expression, which is also increased in distinct sub-populations of cells recovered from human blastocysts. Active expression of HPATs and other HERVH-derived lincRNAs appears necessary for a sustained activity of key pluripotency-regulatory genes since significantly reduction of their expression levels were observed in hESC after small hairpin RNA (shRNA) interference-mediated targeting of HERVH-derived transcripts (Fig.6I; top panel). Notably, siRNA-mediated and CRISPR-targeted interference with the expression of specific *MTTH*-regulated HPAT lincRNAs (*HPAT2; HPAT3;* and *HPAT5*) alters the pluripotent cell fate in human preimplantation embryogenesis and during nuclear reprogramming [17]. Therefore, the results of gene silencing experiments indicate that following activation of HPAT lincRNA’s expression by the core pluripotency transcription factors, the positive feed-back regulatory loop may operate to reinforce the maintenance of the pluripotent state and support a sustained expression of key pluripotency regulators (Fig. 6I). Collectively, the results of our experiments suggest that *MTTH/HPAT* lincRNAs regulatory axis is established in an *HPAT2pos* sub-population of human blastocysts (Fig. 6I). Our analysis indicates that specialization and differentiation of the *HPAT2pos* population, containing putative multi-lineage precursor cells in human embryos, is not limited to the creation of the trophectoderm, primitive endoderm and epiblast lineages. This apparently unique sub-population of human blastocysts may also harbor cells in which the induction of the naïve pluripotency state of hESC occurs (Fig. 6I).

### Single-cell expression profiling of genomic loci encoding HPAT lincRNAs during preimplantation development of human embryos

To further test the hypothesis that *HPAT2pos* sub-populations harbor putative multi-lineage precursor cells, we thought to carry-out single-cell expression profiling analyses during human preimplantation development of TrE-derived lincRNAs, including expression of two HERVH-derived lincRNAs (*HPAT2* and *HPAT3*), which appears markedly enriched in *HPAT2pos* blastocyst cells (Figs. 2 & 3). Significantly, three of five most abundant lincRNAs (*HPAT2; HPAT3;* and *HPAT5*) in human blastocysts were previously implicated in regulation of nuclear reprogramming and pluripotency induction [17]. We reasoned that if the proposed model is valid, the high expression levels of these lincRNAs should be observed in late blastocysts, epiblast cells, and naïve hESCs. To this end, we ascertained the expression profiles of HERV-loci of the human blastocyst HPAT lincRNAs in 153 individual human embryonic cells of the Yan et al. [13] and Xue et al [12] validation data sets (Supplemental Table S1). In striking agreement with the model, expression profiles of HERV-loci of all most abundant lincRNAs (*HPAT21; HPAT15; HPAT5; HPAT2;* and *HPAT3*) in human blastocysts manifest similar patterns of activation in epiblast cells and reach the maximum expression levels in the naïve hESCs (designated in the Fig. 4 as hESCp0 cells). Similar expression patterns of these HERV-loci of HPAT lincRNAs during human preimplantation embryo development were clearly discernable during the analyses of both cells’ sub-population-based profiles plotted as mean expression values (Fig. 4A-E) and expression profiles of 153 individual human embryonic cells at various stages of preimplantation embryogenesis (Fig. 4F-I).

**Figure 4.**
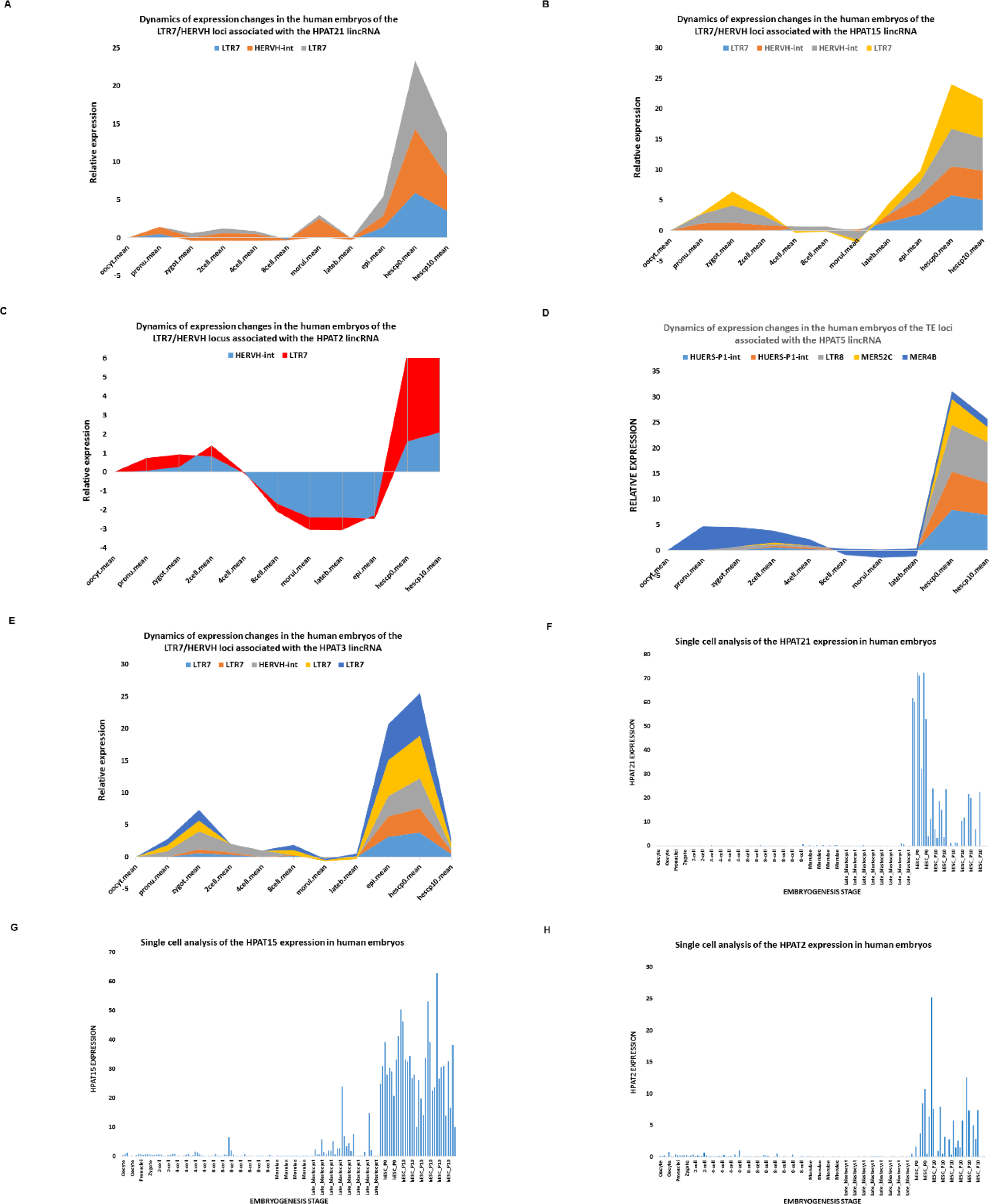
Single-cell analysis of expression patterns of the genetic markers and inducers of the human naive pluripotency state in HPAT lincRNA expression-defined sub-populations during human blastocyst differentiation. Expression changes of designated genetic inducers (A; B) and markers (C; D) of the human naive pluripotency state (see text for details) were evaluated in individual cells comprising the corresponding HPAT expression-defined sub-populations that were recovered from the early-stage and late-stage human blastocysts. The statistical significance of the observed expression changes between sub-populations or within a sub-population during blastocyst differentiation were estimated based on comparisons of the numbers of positive and negative cells using two-tailed Fisher’s exact test. Stars designate sub-populations harboring the significantly different numbers of cells expressing defined genetic markers. P values reported in the Supplemental Table S2. Figure (E) reports the results of HPAT lincRNA expression profiling experiments performed on the OCT4-?PE-GFP cell line after the induction of the naive pluripotency state by overexpression of the *MCRS1, TET1* and *THAP11* genes [18].

We noted that individual human embryonic cells manifesting low expression of TrE-derived lincRNA are detectable throughout the preimplantation embryo development, including the late blastocyst stage. Intriguingly, some TrE-derived lincRNAs manifest clearly distinct expression patterns in naïve hESCs (designated as hESCp0 in Fig. 4) compared with cultured hESCs (designated as hESCp10 in Fig. 4). Individual human embryonic cells with the highest expression of the *HPAT3* lincRNA appear at the late blastocyst stage; they represent all naïve hESCp0 cells, and their prevalence is markedly diminished in population of cultured hESCp10 cells (Fig. 4I). High *HPAT2*-expressing cells appear in the hESC population: they comprise approximately half of the naïve hESCp0 population and continue to persist as a major sub-population in hESCp10 cultures (Fig. 4H). Taken together, these results provide additional strong arguments supporting the validity of our proposed model and indicating that the *HPAT2pos* cells may serve as a putative precursor of naïve hESCs.

### Identification and characterization of putative immortal multi-lineage precursor cells (iMPC) recovered from viable human blastocysts

We reasoned that the multi-lineage precursor cell population may be found among potentially immortal sub-populations of blastocyst cells expressing the human telomerase reverse transcriptase (*TERT*) gene and *TERT* expression may serve as the reliable marker of the putative immortal cells in blastocysts. The importance of telomerase expression for embryogenesis and reproductive potential was clearly established [38]. Furthermore, it was reasonable to expect that *TERT* ^*(+)*^ sub-population may display prominently during the transition from early to late blastocyst stages because all examined human fetal tissues of eight-week of gestation express telomerase [39; 40]. Importantly, it has been shown that telomerase activity and maintenance of telomere stability conferred increased resistance to programmed cell death in human cells [47]. *HPAT* expression-guided reconstruction of human blastocyst differentiation inferred from a single-cell expression profiling identifies a single *HPAT21* ^*(+)*^ *HPAT2* ^*(+)*^ *HPAT15* ^*(+)*^ triple-positive cell expressing the *TERT* gene within the population of the early human blastocysts (Figs. 2F & 5). Significantly, this was the only *TERT*^(+)^ cell observed among 88 early blastocyst cells in our experiments indicating that there is ~1% of *TERT*^(+)^ embryonic cells at this developmental stage (Fig. 2F). The population of *TERT*^(+)^ cells appears to markedly expand during the late blastocyst stage: 56% of *HPAT21* ^*(+)*^ *HPAT2* ^*(+)*^ *HPAT15* ^*(+)*^ triple-positive cells express *TERT* gene in the late blastocyst cell population (Figs. 2G & 5). Similarly, 29% of *HPAT21* ^*(+)*^ *HPAT2* ^*(+)*^ *HPAT15* ^*(−)*^ cells and 3.5% of *HPAT21* ^*(+)*^ *HPAT2* ^*(−)*^ *HPAT15* ^*(+)*^ cells express the *TERT* gene among the late blastocyst cells (Fig. 2G). In striking contrast, all ninety-two *spHPAT21* cells comprising ~38% of the early and late blastocyst cell populations remain *TERT*-negative (Figs. 2F & 2G). These results suggest that *HPAT21* ^*(+)*^ *HPAT2* ^*(+)*^ *HPAT15* ^*(+)*^ and *HPAT21* ^*(+)*^ *HPAT2* ^*(+)*^ *HPAT15* ^*(−)*^ sub-populations designated as the *HPAT2pos* cells are enriched for the putative immortal precursor cell population expressing genetic markers of multiple lineages. Consistent with this hypothesis, we observed that *HPAT2pos* sub-populations are markedly enriched for individual cells expressing genetic markers of the pluripotent state compared with the *spHPAT21* single-positive cells and *dpHPAT15* double-positive cells (Figs. 1–3).

The *HPAT2pos* sub-populations contain a vast majority (16 of 18 cells; 89%) of *TERT* ^*(+)*^ cells identified in human blastocysts in our experiments. Significantly, the functional identities of genes comprising the gene expression signature common for *TERT*^(+)^ cells within the *HPAT2pos* sub-populations appear highly consistent with the potential biological role of these cells in establishing and maintaining a multi-lineage precursor phenotype in human blastocysts (Fig. 5B-D). One of the notable features of the cellular identity of the *TERT* ^(+)^ sub-population is a strikingly distinct pattern of representations of cells expressing specific human endogenous retroviruses (HERV)-derived lincRNAs: a majority (56-100%) of *TERT*^(+)^ cells express the *HPAT21; HPAT2; HPAT5; HPAT15;* and *HPAT3* lincRNAs, whereas there were no *TERT*^(+)^ cells expressing *HPAT4; HPAT6; HPAT7; HPAT9; HPAT14; HPAT17; HPAT20;* and *LINC-ROR* lincRNAs (Figs. 5B-D). These results are highly consistent with the hypothesis that individual HERV-derived lncRNAs may play different biologically-significant roles during the embryonic development [17] and support the idea that *HPAT2pos* sub-populations of human blastocysts contain putative immortal multi-lineage precursor cells (iMPCs).

**Fig. 5.**
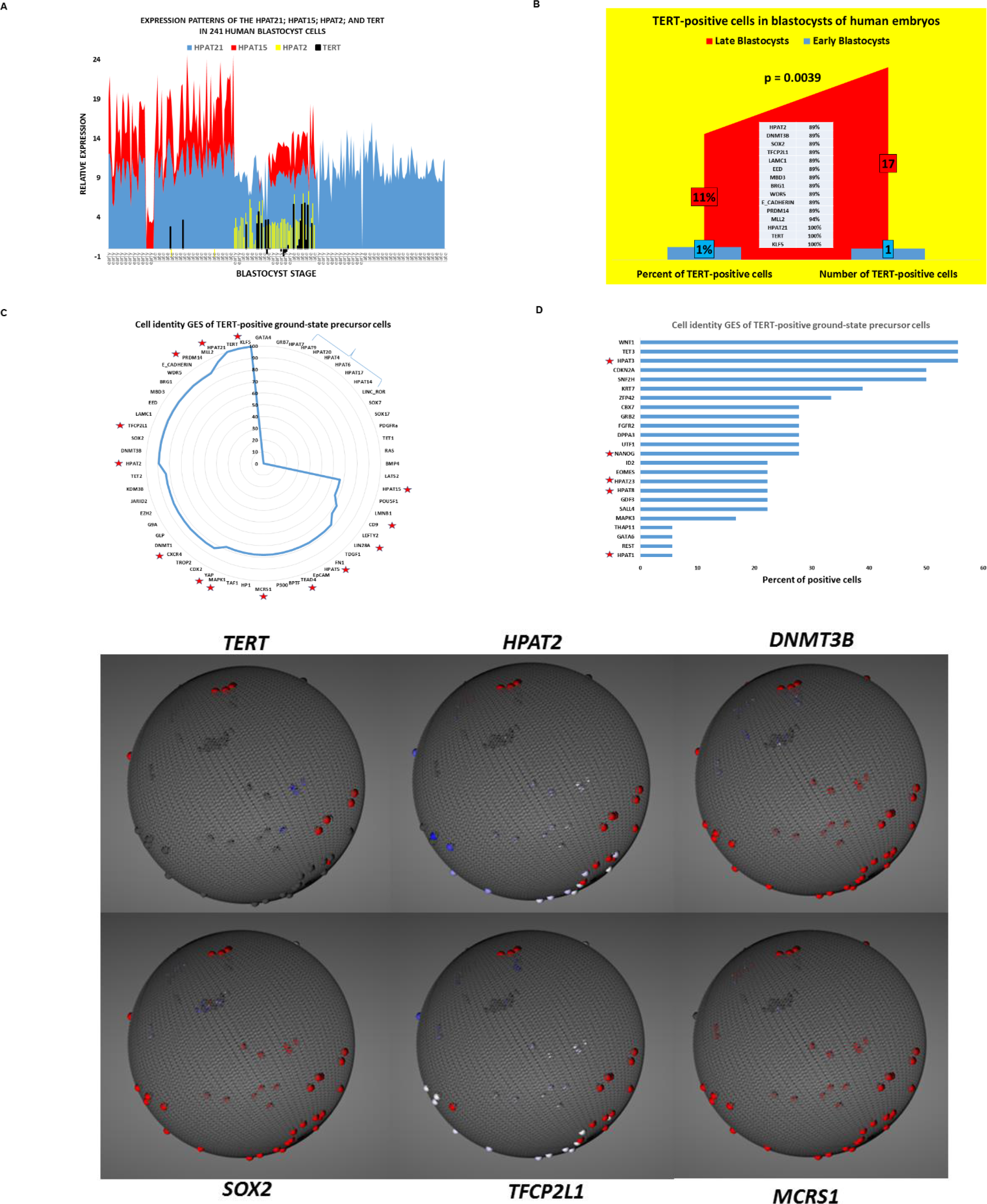
Identification and characterization of putative immortal multi-lineage precursor cells created during human blastocyst differentiation. **(A)** Expression profiles of the human telomerase reverse transcriptase *(TERT)* gene transcript and *HPAT21; HPAT15;* and *HAPT2* lincRNAs in human blastocyst cells. A color-coded stacked area plot depicts relative expression values of the *TERT* gene transcript and designated HPAT lincRNAs in individual blastocyst cells. Note that a vast majority (89%) of *TERT* ^(+)^ cells emerged within the HPAT21 ^(+)^ HPAT15 ^(+)^ HPAT2 ^(+)^ & HPAT21 ^(+)^ HPAT15 ^(-)^ HPAT2 ^(+)^ sub-populations of human blastocyst cells. **(B)** Marked expansion of *TERT* ^(+)^ cells during human blastocyst differentiation. Note that there is only a single *TERT* ^(+)^ cell was detected in the population of the early-stage blastocyst cells (~1%) whereas the *TERT* gene expression was detected in a relatively large sub-population (11%) of the late-stage blastocyst cells. The inset lists genes that are expressed in a vast majority (89% to 100%) of *TERT* ^*(+)*^ cells recovered from human blastocysts. **(C)** Cell identity gene expression signature (GES) of *TERT* ^*(+)*^ cells recovered from human blastocysts. The list includes genes that are not expressed in *TERT* ^*(+)*^ cells and genes that are expressed in at least two-third of *TERT* ^*(+)*^ cells (from 67% to 100% of telomerase-positive cells). Stars in (C) and (D) designate genes that were identified as pluripotency regulators, inducers, or markers of the pluripotent state, including eight HPAT lincRNAs that are expressed in *TERT* ^*(+)*^ cells. Right brace in (C) designates eight lincRNAs expression of which appears repressed *TERT* ^*(+)*^ cells. Positions of the blue line inside the circle reflect the percentage of cells within the TERT ^(+)^ population that express transcripts encoded by the corresponding genes. **(D)** List of genes that are expressed in 6%-56% of *TERT* ^*(+)*^ cells recovered from human blastocysts. Bar heights correspond to the percentage of *TERT* ^*(+)*^ cells expressing designated genes. **(E)** Visualization of *TERT* ^*(+)*^ putative immortal ground-state pluripotency precursor cells using the 3D reconstructed blastocyst model that was utilized to design the first web-based online tool to study early cell fate decisions in the human blastocyst, which is available online at http://web.stanford.edu/~sunilpai/HumanBlastocystViewer.html (Firefox/Chrome compatible). A single frame of the 3D blastocyst sphere is shown that captured ten TERT ^(+)^ human late blastocyst cells. Expression patterns of other genetic markers in individual human blastocyst cells located within the same field of the 3D reconstructed blastocyst sphere are shown. The extended report of this analysis is available in the Supplemental Movie.

### Single cell analysis of *TERT* expression dynamics in human preimplantation embryos revealed a timeline of emergence and sustained presence of putative iMPCs

We carried out a series of follow-up analyses to identify human embryonic cells resembling the putative iMPCs among individual cells and distinct lineages which were identified and characterized in a recently reported independent data set of single cell expression profiles of 1,529 embryonic cells recovered from 88 human embryos at different stages of preimplantation embryonic development [34]. To this end, we evaluated *TERT* mRNA expression patterns in each of the 1,529 individual human embryonic cells that were recovered at different stages of preimplantation embryogenesis and assigned to distinct lineages based on their gene expression (Supplemental Figure S4). These analyses demonstrate that *TERT*^(+)^ cells could be observed among all lineages and are readily detectable as early as at E3-E4 stages followed by the expansion during the E5 stage coincidently with blastocyst formation (Supplemental Figure S4). We found that E5 pre-lineage cells manifest significantly increased levels of *TERT* mRNA expression compared to EPI, PE, and TE lineages (Supplemental Figure S4). Interestingly, the second peak of *TERT* activity was observed at the E6 in all three major lineages created during human preimplantation embryonic development. We observed that the E5.early cell population has the highest level of *TERT* mRNA expression and contains the largest percentage of *TERT*^(+)^ cells compared to other stages of human preimplantation embryonic development (Supplemental Figure S4). Therefore, based on the results of analyses of *TERT*^(+)^ cells in human preimplantation embryos we built a quantitative model of *TERT*^(+)^ cells’ dynamics in an individual human embryo demonstrating that:

i) *TERT*^(+)^ cells pervade all lineages and are readily detectable throughout the E3-E7 stages of embryonic development (Supplemental Figure S4);
ii) E5 pre-lineage population expresses the highest level of *TERT* mRNA compared to E5.EPI, E5.PE, and E5.TE lineages;
iii) E5.early cells appear most enriched for telomerase-positive cells and express the highest level of telomerase mRNA (Supplemental Figure S4).

Petropoulos et al. [34] reported that cells during human embryonic development acquire an intermediate state of co-expression of lineage-specific genes, followed by a concurrent establishment of the trophectoderm (TE), epiblast (EPI), and primitive endoderm (PE) lineages coincidently with blastocyst formation. They concluded that segregation of all three lineages in human embryos occurs simultaneously and coincides with blastocyst formation at E5. During the early E5 stage, embryonic cells that had activated TE genes also maintain the expression of EPI genes, consistent with the hypothesis of an intermediate stage of co-expression of genetic markers of multiple lineages [34]. We reasoned that cells within the cell population co-expressing genetic markers of multiple lineages and termed E5 pre-lineage may be phenotypically similar to identified herein putative immortal multi-lineage precursor cells (iMPC). Consistent with this line of reasoning, we observed that the E5 pre-lineage population harbors a sub-population of *TERT*^(+)^ cells resembling the putative iMPCs (Fig. 6).

**Figure 6.**
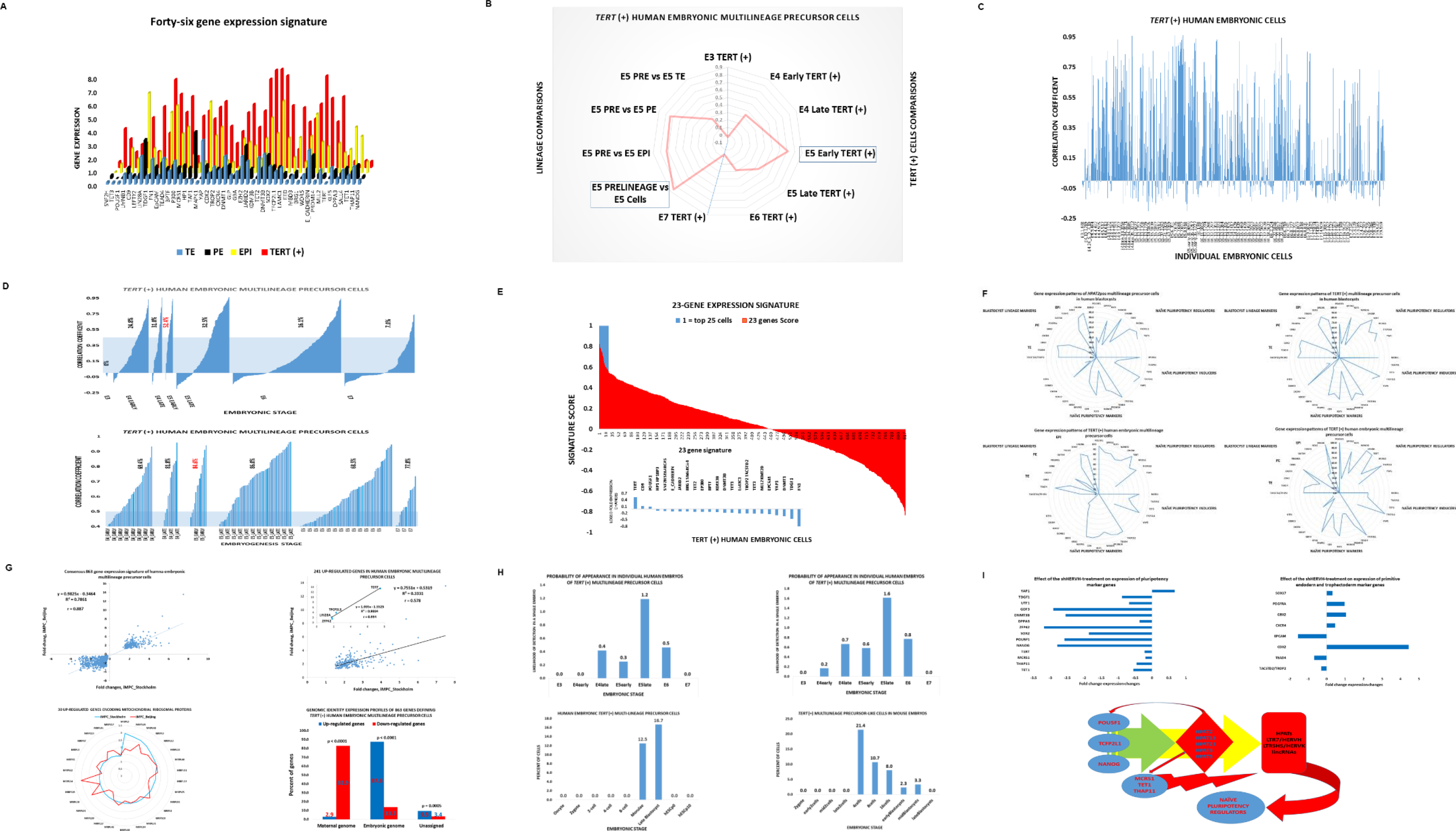
Identification of *TERT* ^(+)^ cells resembling the putative immortal multi-lineage precursor cells (iMPCs) in independent validation sets of 1,708 individual human embryonic cells recovered from more than 100 human embryos at distinct stages of preimplantation human embryonic development. **(A)** Forty-six gene expression signature of the iMPC population identified in the single-cell gene expression profiling experiments of 241 human blastocyst cells recovered from 32 human embryos (discovery data set; Figs. 1–5) was utilized to identify the iMPC-resembling cells among the 1,529 individual embryonic cells recovered from 88 human embryos at distinct stages of preimplantation embryonic development (validation data set 1; Supplemental Table S1 and ref. 34). **(B)** A graphical summary of the population-based correlation score analyses. Note that E5.Pre-lineage cells and E5.Early *TERT* ^*(+)*^ cells manifest the most significant enrichment for the iMPC-like cells in the validation data set 1. Correlation patterns of the forty-six gene iMPC signature defined by gene expression ratios of *TERT* ^*(+)*^ cells versus EPI cells (discovery data set) and E5.Pre-lineage cells versus E5.EPI cells (validation data set 1). High positive values of the correlation score indicate the resemblance to the iMPC gene expression profile. Similarly, correlation scores were calculated for the 46-gene iMPC signature defined by the gene expression ratios of *TERT* ^*(+)*^ cells versus EPI/PE/TE cells (discovery data set) and E5.Pre-lineage cells versus E5.EPI/PE/TE cells (validation data set 1); for the 46-gene iMPC signature defined by the gene expression ratios of *TERT* ^(+)^ cells versus EPI/PE/TE cells (discovery data set) and E5.Early *TERT* ^*(+)*^ cells versus E5 cells (validation data set 1); and for the forty-six gene iMPC signature defined by the gene expression ratios of *TERT* ^*(+)*^ cells versus EPI/PE/TE cells (discovery data set) and E5.Early *TERT* ^*(+)*^ cells versus E5 cells (validation data set 1). Finally, correlation score analysis of the 46-gene iMPC signature defined by the gene expression ratios of the *TERT* ^*(+)*^ cells versus EPI/PE/TE cells (discovery data set) and E3. *TERT* ^*(+)*^ cells versus E3 cells (validation data set 1). Note that in this instance no significant correlation was observed. **(C)** Correlation score patterns of the forty-six gene iMPC signature defined by the gene expression ratios of the *TERT* ^*(+)*^ cells versus EPI cells (discovery data set), which were assessed in 819 individual *TERT* ^*(+)*^ cells identified in the validation data set 1. Cells were placed in the ascending order of the individual identification numbers from E3 (left) to E7 (right) stages of embryonic development. **(D)** Correlation score patterns of the forty-six gene iMPC signature defined by the gene expression ratios of the *TERT* ^*(+)*^ cells versus EPI cells (discovery data set) which were assessed in 819 individual *TERT* ^*(+)*^ cells identified in the validation data set 1. Cells were sorted in ascending order of correlation scores within corresponding developmental stages (top panel). Bottom panel shows the zoom-in view of the single-cell analysis of a subset of cells of the validation data set 1, which were segregated at a cut-off value of correlation score 0.4. Percentage values are the percent of individual cells within a population with the correlation score > 0.5. Using this approach, a total of 158 *TERT* ^*(+)*^ human embryonic cells manifesting correlation scores > 0.5 were identified in the validation set 1. **(E)** Twenty-three gene signature comprising a sub-set of genes of the 46-gene signatures that manifest significantly different expression levels in 158 *TERT* ^*(+)*^ cells (r > 0.5) compared to 661 *TERT* ^*(+)*^ cells (r < 0.5) in the validation data set 1. Using the gene expression profile of the 23-gene signature, correlation scores were calculated for 819 *TERT* ^*(+)*^ cells of the validation data set 1 (red colored bars). Blue bars designate top-scoring twenty-five *TERT* ^*(+)*^ cells manifesting correlation scores r > 0.55. **(F)** Similarity patterns of gene expression signatures (GES) defining putative immortal multi-lineage precursor cells identified in the Durruthy-Durruthy et al. discovery data set (top two figures), in the Petropolous et al. validation data set 1 (bottom left figure), and in the Yan et al. validation data set 2 (bottom right figure). The top two figures show GES of the *HPATpos* cells (left) and *TERT* ^*(+)*^ cells (right) identified in the Stanford discovery data set. The bottom left figure shows GES of the top-scoring *TERT* ^*(+)*^ cells identified in the Petropolous et al. validation data set 1 (Supplemental Table S1) based on the correlation scores of the 23-gene signature. The bottom right figure shows GES of the top-scoring *TERT* ^*(+)*^ cells identified in the Yan et al. validation data set 2 (Supplemental Table S1) based on the correlation scores of the 23-gene signature. The genes comprising the GES were identified in the single cell expression profiling experiments of viable human blastocysts (the Stanford discovery data set). The gene names listed around the circles were grouped into relevant functional categories and placed in the same orders in all four figures. The numerical values corresponding to each gene indicate the percentage of positive cells which were identified using defined expression thresholds within the corresponding populations (gene expression values above null in the discovery data set and median expression values for non-iMPC-resembling cells in validation data sets). **(G)** Consensus 863-gene expression signature of the *TERT* ^*(+)*^ human embryonic multi-lineage precursor cells (iMPC). A total of 863 genes (241 up-regulated and 622 down-regulated genes) manifesting highly concordant gene expression profiles (r = 0.887; top left figure) in the *TERT* ^*(+)*^ human embryonic multi-lineage precursor cells were identified using two independent validation data sets (Petropolous et al. validation data set 1 and Yan et al. validation data set 2). Concordant expression profiles of 241 up-regulated genes of the consensus gene expression signature of iMPC are shown in the top right figure. The inset highlights concordant expression patterns of four key pluripotency regulatory genes (TERT; TCFP2L1; LIN28A; and ZFP42). Bottom left figure shows expression patterns of 30 genes encoding mitochondrial ribosomal proteins, expression of which is significantly increased in both populations of TERT ^(+)^ human embryonic iMPC identified in two independent validation data sets. Bottom right figure documents distinct genomic origins of 241 up-regulated (blue bars) and 622 down-regulated (red bars) genes comprising the consensus gene expression signature of the TERT ^(+)^ human embryonic iMPC. Note that transcription of 87.6% up-regulated genes originates from the embryonic genome, while transcriptional origins of 82.9% down-regulated genes were assigned to the maternal genome. The assignments of genomic origins of 863 genes were performed based on single cell gene expression profiling of twenty-six human embryonic cells reported in the validation data set 4 (Supplemental Table S1). **(H)** Timelines of creation of *TERT* ^*(+)*^ embryonic immortal multi-lineage precursor cells (iMPC) during the preimplantation embryogenesis in humans and mouse. The top left & right figures show the human embryogenesis timelines based on the assessments of the top-scoring *TERT* ^*(+)*^ cells identified in the Petropolous et al. validation data set 1 (Supplemental Table S1), which were selected based on the correlation scores of the 23-gene signature for top 3% *TERT* ^*(+)*^ cells (top left figure) and top 5% *TERT* ^*(+)*^ cells (top right figure). The bottom left figure reports the human embryogenesis timeline based on the top-scoring *TERT* ^*(+)*^ cells identified in the Yan et al. validation data set 2 (Supplemental Table S1), which were selected based on the correlation scores of the 23-gene signature for 8% of TERT ^(+)^ cells. The estimated numbers of the likelihood of appearance of the human iMPC-resembling cells in a single human embryo were calculated based on the assumption that the numbers of human embryonic cells were 8; 16; 32; 64; and 128 at the E3; E4; E5; E6; and E7 stages, respectively. The human iMPC-like cells were defined as *TERT* ^*(+)*^ cells comprising the top 3% (top left figure) and top 5% (top right figure) of the 819 TERT ^(+)^ cells defined based on the 23-gene signature correlation scores in the validation data set 1 (top two figures). The proportions of cells in the E4.early versus E4.late and E5.early versus E5.late populations within the validation set 1 were estimated from the data reported in [34]. The human iMPC-like cells reported in the bottom left figure were defined as *TERT* ^*(+)*^ cells comprising the top 8% (bottom left figure) of cells defined based on the 23-gene signature correlation scores in the validation data set 2. Bottom right figure shows the mouse embryogenesis timeline of creation of iMPC-like cells based on the top-scoring *TERT* ^*(+)*^ cells identified in the mouse preimplantation embryogenesis data set (Supplemental Table S1) comprising 259 individual embryonic cells. The top-scoring 13.9% of *TERT* ^*(+)*^ mouse embryonic cells were identified based on the correlation scores of the human iMPC consensus 863-gene expression signature (Fig. 6G). **(I)** A model of principal molecular events contributing to regulation of the naïve pluripotency induction in vivo during transition from *TERT* ^*(+)*^ iMPC to pluripotent epiblast and hESC in human embryos. The first wave of increased expression of 5 most abundant in human blastocysts HPAT lincRNAs (highlighted by the green arrow) is triggered by the master pluripotency transcription factors (POU5F1/OCT4; TCFP2L1/LBP9; NANOG). The positive feed-back regulatory loop mediated by the activity of HPAT lincRNAs (red arrows) increases expression of master pluripotency transcription factors and naïve pluripotency inducers (MCRS1; TET1; THAP11), which triggers the second wave of increased expression of a multitude of TrE-derived lincRNAs (HPATs; LTR7/HERVH; and LTR5HS/HERVK families). Increased activities of TrE-derived lincRNAs make a critical contribution to the embryonic cells’ chromatin remodeling by enabling transitions to thermodynamically-stable triple-stranded state of double helix and facilitating targeted delivery of POU5F1/OCT4 & Mediator proteins to thousands of genomic loci. A second wave of the positive feed-back regulatory loop mediated by the activities of TrE-derived lincRNAs increases expression of key naive pluripotency regulators. Top two figures show the graphical summary of the effects of shRNA-mediated targeted knockdown of LTR7/HERVH lincRNAs in hESC inducing statistically significant changes in expression of genes encoding naive pluripotency regulators (top left figure) and genetic markers of TE and PE lineages (top right figure). Experimental evidence supporting the model are reported and discussed in the text.

To detect cells resembling the putative iMPCs among the 1,529 individual embryonic cells reported in the Petropolous et al. study [34], we carried out correlation analyses of expression profiles of forty-six genes distinguishing iMPCs from other lineages (Figs. 5 & 6; Supplemental Table S3). In these experiments, the expression profiles of forty-six genes comprising candidate cell identity genes of iMPCs were identified in individual cells and different lineages in the Petropolous et al. data set [34] and compared to the iMPC expression profile of the Stanford data set [18]. Cells were defined as iMPC-resembling if their correlation coefficient of gene expression profiles exceeds 0.5. These analyses revealed the consistent presence of iMPC-resembling cells among human embryonic cells characterized in the Petropolous et al. data set (Fig. 6). Importantly, the iMPC-resembling cells are represented most prominently among E5 pre-lineage cells and *TERT*^(+)^ E5.early cells of the Petropolous et al. study (Fig. 6B). In striking contrast, the population of *TERT*^(+)^ E3 cells does not appear to contain iMPC-like cells (Fig. 6B). This conclusion remains valid when observations were made employing the population-based approach by comparing either the lineage-defined or embryonic stage-defined populations (Fig. 6).

We thought to extend these population-based observations by performing the analyses of individual human embryonic cells. To this end, we selected all 819 *TERT*^(+)^ cells identified in the Petropolous et al. data set and compared their individual gene expression profiles to the 46-gene iMPC expression signature (Figs. 6C & 6D). Consistently with the conclusions made based on the analyses of the embryonic lineage-defined and embryonic stage-defined populations (Fig. 6B), there were no iMPC-like individual cells detectable at the E3 stage (Fig. 6D). We observed that individual iMPC-resembling cells emerged among human embryonic cells at the E4.early stage and persisted throughout E4-E7 stages of human embryonic development (Fig. 6D). The largest percentage of the iMPC-resembling cells was observed among the *TERT*^(+)^ E5.early cells (Fig. 6D). Using this strategy, we identified *158 TERT* ^*(+)*^ human embryonic cells in the Petropolous et al. data set, which recapitulate the 46-gene expression profile of the iMPCs (Figs. 6C & 6D). Furthermore, our analyses identify the E5 pre-lineage cells and the E5.early *TERT* ^*(+)*^ cells as sub-populations of human embryonic cells that appears most enriched for iMPC-like cells. Therefore, it seems reasonable to conclude that these analyses demonstrate a consistent presence of iMPC-resembling cells in human preimplantation embryos.

### Genome-wide screens of single-cell expression profiles of embryonic cells identifies consensus gene expression signatures of the iMPCs in human and mouse embryos

To further test the validity of our hypothesis that iMPCs are created during human preimplantation embryogenesis, we carried-out several additional follow-up analytical experiments. We performed the formal statistical analysis to determine whether the expression levels of the forty six genes comprising the iMPC gene expression signature are statistically different in 158 *TERT* ^*(+)*^ iMPC-like cells compared with 661 *TERT* ^*(+)*^ cells not resembling the iMPC of the Petropolous et al. validation data set. This analysis identifies twenty three genes (50%) manifesting statistically distinct expression levels in *TERT* ^*(+)*^ iMPC-like cells (Fig. 6E). We thought to utilize this 23-gene expression signature for more stringent definition of iMPC-like cells in independent validation data sets of single-cell expression profiling of human preimplantation embryos [13; 34]. We considered that the cut-off number of top-scoring *TERT* ^*(+)*^ cells should be sufficient to estimate that at least one iMPC-like cell would emerge in individual human embryos at any embryogenesis stage prior to the lineage segregation. Therefore, in subsequent validation analyses we thought to restrict the selection of putative iMPC-like cells in a data set to the minimum number of *TERT* ^*(+)*^ cells displaying the highest 23-gene signature correlation scores, provided the above criterion is satisfied. Using the expression profile of 23-gene signature (Fig. 6E; Supplemental Table S3) as a tool for more stringent definition of iMPC-resembling cells, we identified top-scoring *TERT* ^*(+)*^ iMPC-like embryonic cells in two independent validation data sets comprising 1,529 and 124 individual human embryonic cells and designated as validation data set 1 and 2, respectively (Supplemental Table S1; refs. 34 & 13).

Next, the similarity patterns of gene expression signatures (GES) defining putative immortal multi-lineage precursor cells were ascertained by comparisons of three populations of iMPCs which were independently identified in the Stanford discovery data set (Fig. 6F; top two figures), in the Petropolous et al. validation data set 1 (Fig. 6F; bottom left figure), and in the Yan et al. validation data set 2 (Fig. 6F; bottom right figure). The genes comprising the iMPC GES were identified in the single cell expression profiling experiments of viable human blastocysts (Stanford discovery data set; Figs. 1–5), grouped into relevant functional categories and placed in the same orders in all four images shown in the Fig. 6F. The top two figures show the iMPC GES profiles of the *HPATpos* cells (left) and *TERT* ^*(+)*^ cells (right) identified in the Stanford discovery data set. The bottom left figure shows the iMPC GES profile of the top-scoring *TERT* ^*(+)*^ cells identified in the Petropolous et al. validation data set 1 (Supplemental Table S1) based on the correlation scores of the 23-gene signature. The bottom right figure shows the iMPC gene expression profile of the top-scoring *TERT* ^*(+)*^ cells identified in the Yan et al. validation data set 2 (Supplemental Table S1) based on the correlation scores of the 23-gene signature. The numerical values corresponding to each gene indicate the percentage of positive cells which were identified using defined expression thresholds within the corresponding populations and plotted to observe the images of signature patterns (Fig. 6F). We conclude that gene expression profiles’ visualization analysis demonstrates clearly discernable similarity patterns of GES in independently-defined populations of *TERT* ^*(+)*^ human iMPCs.

Identification of the iMPC-like cells in independent validation data sets of human preimplantation embryogenesis afforded the opportunity to utilize genome-wide gene expression data of individual embryonic iMPC-like cells for definition of a consensus gene expression signature of *TERT* ^*(+)*^ iMPC in human embryos. We accomplished this task by identifying differentially expressed genes distinguishing the iMPC-like cells in the Petropolous et al. and Yan et al. data sets and selecting the genes manifesting the concordant statistically significant up- and down-regulation in both data sets. This approach identifies the 863-gene consensus expression signature of *TERT* ^*(+)*^ iMPC of human embryos comprising 241 up-regulated and 622 down-regulated genes (Fig. 6G). Notably, the genomic origin of a majority (82.9%) of down-regulated genes of the consensus *TERT* ^*(+)*^ iMPC expression signature could be traced to the maternal genome (Fig. 6G; red bars in the bottom right image), suggesting that the efficient transcriptional repression of maternally-expressed genes represents one of the key features of the iMPC population. In striking contrast, the genomic origin of a vast majority (87.6%) of up-regulated genes could be traced to the embryonic genome (Fig. 6G; blue bars in the bottom right image), indicating that genes transcriptionally active in the embryonic genome make a significant contribution to the creation of the *TERT* ^*(+)*^ iMPC in human embryos.

We thought to employ the 23-gene expression signature of *TERT* ^*(+)*^ iMPC-like cells of human embryos to identify the putative iMPC-like cells during the mouse preimplantation embryogenesis [45]. Analysis of gene expression profiles of 259 mouse embryonic cells (mouse embryo validation data set; Supplemental Table S1) identifies top-scoring *TERT* ^*(+)*^ iMPC-like mouse embryonic cells comprising 5% of mouse embryonic cells. Using genome-wide statistical screens of single-cell gene expression profiling of embryonic cells, we were able to identify individual iMPC-like cells in a data set which were assigned to the particular stages of human and mouse embryonic development. Using these data and reported numbers of analyzed embryonic cells assigned to the corresponding embryogenesis stage in a data set, we estimated the likelihood of emergence and expected creation timelines of the *TERT* ^*(+)*^ iMPC in individual human and mouse embryos (Fig. 6H). We observed a sustained presence of the *TERT* ^*(+)*^ iMPC-resembling cells during the E4-E6 stages of human preimplantation embryonic development in the Petropolous et al. validation data set 1 (Fig. 6H; top two images). In the Yan et al. validation dataset 2, the creation timelines of the *TERT* ^*(+)*^ iMPC in human embryos were defined as the morulae and blastocyst stages (Fig. 6H; the bottom left image). In contrast to human embryos, the *TERT* ^*(+)*^ iMPC-like cells appear to emerge as early as the 4-cell and 8-cell stages of the mouse embryonic development (Fig. 6H; the bottom right image). Therefore, iMPC-like cells seem to emerge earlier during mouse preimplantation embryogenesis compared with human embryos (Fig. 6H; the bottom right image). These observations are consistent with the model of embryonic development that was proposed based on the mouse preimplantation embryogenesis studies, where the TE and ICM fate is initiated in a positional and cell polarization-dependent manner within the morulae [41] and is followed by a progressive maturation of the EPI and PE lineages in the blastocyst [42]. Consistent with this line of reasoning, human morulae compaction occurs at the 16-cell stage [43] whereas mouse morulae compaction takes place at the 8-cell stage, explaining a delay in lineage segregation observed in the human embryos compared with the mouse [34].

The results of extensive validation analyses using single cell expression profiles of the 1,708 individual embryonic cells recovered from more than 100 human embryos and 259 individual mouse embryonic cells at different stages of preimplantation embryonic development appear to support the validity of the model that during embryonic development the *TERT* ^*(+)*^ iMPC population is created. Based on the previously published studies [3; 8; 15; 17; 18; 35] and experimental evidence reported in this contribution, we propose a model of principal molecular and cellular events contributing to regulation of the naïve pluripotency induction *in vivo* during transition from *TERT* ^*(+)*^ iMPC to pluripotent epiblast cells and hESCs in human embryos (Fig. 6I). The key elements of this two-wave embryogenesis progression model highlight the central regulatory role of the TrE-derived lincRNAs and the *MTTH/HPAT* lincRNAs axis in promoting a genome-wide transition to the embryonic chromatin state leading to creation of ground-state naïve hESC population (Fig. 6I).

Gene ontology analysis strongly supports definition of *TERT* ^*(+)*^ iMPCs as immature multi-lineage precursors lacking gene expression features of specialized cells (Supplemental Figure S5). Consistent with the proposed role of HPAT lincRNAs in the biogenesis of iMPCs, the most statistically enriched biological processes among genes up-regulated in iMPCs are non-coding RNA metabolism and non-coding RNA processing (Supplemental Fig. S5). Other significantly enriched biological processes among genes up-regulated in iMPCs are translation, tRNA metabolic processes, establishment of protein localization, protein transports, ribonucleoprotein complex biogenesis (Supplemental Fig. S5). Therefore, gene ontology analysis appears to indicate that gene expression networks of iMPCs are devoted to establishing key fundamental building blocks of an essential intracellular infrastructure that is required for proper biological functions of all cells. We noted that genes encoding mitochondrial ribosomal proteins represent one of the largest families of transcripts up-regulated in iMPCs (Fig. 6G; bottom left image). Since mRNAs for all 13 proteins that are necessary to assemble mitochondria are encoded by mitochondrial DNA and translated on mitochondrial ribosomes, these data suggest that activation of genomic networks supporting the energy-producing infrastructure in a cell is one of the key features of iMPCs. Furthermore, mitochondrial ribosomal proteins were implicated in ribosomal RNA-mediated protein folding by facilitating the release of proteins in a folding competent state [49].

In striking contrast, biological processes that are significantly enriched among genes down-regulated in iMPCs appear linked with cellular differentiation and fine specialization of differentiated cells (Supplemental Figure S5). They were defined by gene ontology analysis as processes of cell adhesion, biological adhesion, cell motion, cell motility, localization of cells, regulation of response to external stimulus, cell migration, response to wounding, and gland development. Collectively, these results are highly consistent with observations that iMPCs represent a transitory cell type during embryogenesis (Fig. 6H) that differentiate into trophectoderm, primitive endoderm and pluripotent epiblast lineages.

## Discussion

The novel elements of the experimental approach implemented in this study are based on the isolation of viable human blastocysts and separation of the human blastocyst cells into early and late blastocyst cell populations based on strict morphological criteria defined by the classic embryology. We believe that this approach provides a more stringent basis to control human embryonic cells’ viability. High level of certainty that we analyzed the viable human embryonic cells should mitigate potential limitations of previous studies utilizing the single cell expression profiling approaches of the analysis of human preimplantation embryogenesis and relying solely on segregation of individual cells based on the time after fertilization. One of the obvious flaws of this timeline-based approach is that integration of the single cell analyses from different human embryos would inevitably generate the developmentally heterogeneous sub-populations simply because the lack of synchronization of embryonic development among individual human embryos.

Application of this experimental approach in combination with the state-of-the-art single cell gene expression analysis and spatiotemporal reconstruction of human blastocyst differentiation facilitated the development of a three-dimensional model of the inner cell mass and trophectoderm in which individual cells were faithfully mapped into distinct expression domains [18]. Using this approach, we were able to extract and describe the key novel mechanistic information emanated by the single-cell expression analyses of protein-coding genes [18]. Specifically, these experiments revealed a previously uncharacterized role for *MCRS1, TET1,* and *THAP11* in epiblast formation and their ability to induce naive pluripotency *in vitro.*

Remarkably, the single cell expression profiling analysis of human blastocyst differentiation based solely on the TrE-derived lincRNAs expression profiles facilitated a series of novel observations potentially leading to significant conceptual advances in our understanding of genetic, molecular, and cellular mechanisms of human preimplantation embryonic development. Identification and characterization of putative iMPCs highlighted a key role of *MTTH/HPAT* lincRNAs regulatory axis in a two-wave progression sequence of human embryonic development (Fig. 6I). Significantly, sequence conservation analyses of HPAT lincRNAs, expression profiling of which facilitated identification of putative iMPCs, indicated that they represent human-specific (unique to human) regulatory molecules (data not shown). A hallmark feature of human-specific integration sites of stem cell-associated retroviral sequences, including specific HPAT lincRNAs, is deletions of ancestral DNA [35].

It has been shown that activation of specific HERV loci, including *HPAT3* and *HPAT4* lincRNAs, occurs in human zygotes following *in vitro* fertilization of mature oocytes [47]. Gene expression signature analysis of early-stage human preimplantation embryogenesis revealed that "premature" activation of HERVH expression is associated with increased likelihood of failure to produce viable blastocysts by human fertilized oocytes [47]. These observations support the hypothesis that premature activation of HERV expression in early-stage human embryos may reflect failed attempts to create iMPCs. Collectively, this analysis suggests that premature activation of HERV expression in human embryonic cells may cause cell cycle arrest and loss of cell viability, leading to exceedingly high level of cellular extinction events.

Observations supporting the hypothesis that putative iMPCs are created during the human blastocysts differentiation were facilitated by the convergence of two synergistic analytical advances: identification and initial structural-functional characterization of genomic loci encoding HPAT lincRNAs and technical implementations of the idea of performing a single-cell analysis of viable human blastocyst differentiation by segregating early-stage and late-stage blastocyst cells. Segregation and transcriptional profiling of individual viable human blastocyst cells separated in time by just few critical hours of embryonic development enabled to dissect the dynamics of human blastocyst differentiation with unprecedented precision which was validated by the accurate classification of distinct blastocyst lineages based on the expression profiles of just three HPAT lincRNAs, namely *HPAT21; HPAT15;* and *HPAT2.*

Discovery of HPAT lincRNAs was facilitated by application of a hybrid RNA sequencing technique enabling the identification of more than 2,000 new lincRNA transcript isoforms, of which 146 were specifically expressed in pluripotent hESCs [36]. Follow-up experiments revealed that 23 genomic loci encoding TrE-derived transcripts and termed *HPAT1-HPAT23* lincRNAs are most abundantly and specifically expressed in human pluripotent cells [17]. It has been demonstrated that three specific TrE-derived HPAT lincRNAs *(HPAT2, HPAT3* and *HPAT5)* may modulate cell fate in human preimplantation embryo development and during nuclear reprogramming [17]. Our experiments identified five most abundant HPATs *(HPAT21; HPAT15; HPAT5; HPAT2;* and *HPAT3)* expression of which was observed most frequently in human blastocyst cells. Expression of the *HPAT21* lincRNA was detected in 92% and 100% of cells recovered from early and late stage-developing blastocysts, respectively, suggesting that this lincRNA could be defined as the pan marker of human blastocyst cells.

The conclusive unequivocal validation of the hypothesis that during human blastocyst differentiation the unique sub-populations of putative iMPCs are created, which fuels the emergence of the trophectoderm, primitive endoderm and epiblast lineages, and induction of the naïve hESCs, may prove challenging since it would require the direct isolation and recovery of these unique cells from the human blastocysts and their comprehensive molecular and functional characterization. However, our observations that expression of HPAT lincRNAs implicated in human blastocyst differentiation is markedly up-regulated in human cells acquiring the genetically-induced naïve pluripotency state suggest that it might be possible to design the experimental protocols for genetic engineering of the iMPC-resembling cells. It will be of interest to investigate the developmental potential and translational utility of iMPCs for regenerative medicine.

Collectively, our observations demonstrate that during preimplantation embryogenesis the iMPCs are created and suggest that subsequent differentiation and specialization of these cells contributes to the emergence of the trophectoderm, primitive endoderm and epiblast lineages and induction of naïve hESCs. Our conclusions are based on single cell expression analyses of 241 individual cells recovered from 32 human embryos at early and late stages of blastocyst differentiation. We validated our principal findings and the key predictions of our model by the *in silico* follow-up analyses on 1,708 individual embryonic cells recovered from more than 100 human embryos at different stages of preimplantation development. Our findings are in agreement with recent reports suggesting that single-cell gene expression profiling of cellular differentiation events in human preimplantation embryos and created *in vivo* naive pluripotent stem cells represents the most promising approach to reliably benchmark ground state pluripotency regulatory networks in human cells [17; 18; 37].

## Experimental Procedures

### Human blastocyst retrieval and disaggregation into single cells

Frozen human blastocysts at day 5 and day 6 post fertilization of preimplantation development were thawed according to Quinn’s advantage thaw kit (CooperSurgical, Trumbull, CT) as previously described (Wong et al 2010). Briefly, cryocontainers were removed from liquid nitrogen and exposed to air for 10 sec before incubating in a water bath at 37 °C until thawed. Next, embryos were transferred to 0.5 M and 0.2 M sucrose solution for 10 min each and washed in diluent solution for 10 min at room temperature (RT). Blastocysts were then incubated in Quinn’s Advantage Blastocyst Medium supplemented with 10% serum protein substitute under mineral oil at 37 °C with 6% CO2, 5% O2 and 89% N2, standard human embryo culture conditions for 5 or 12 hours respectively. Only morphological normal looking blastocysts with Gardner grading 1-3AABB (early blastocyst) and 4-6AABB (late blastocyst) (Gardner et al., 2000) were used for the further analysis. Blastocysts were treated with acidic Tyrode’s solution and washed in Dulbecco’s PBS (DPBS) substituted with 1 mM EDTA and 0.5% BSA. After incubation in 0.005% trypsin-EDTA for 15 mins blastocyst were washed in DPBS and collected for the following C1-analysis.

The *MTTH* model of induced human naïve pluripotent cells was established and analyzed using the OCT4-ΔPE-GFP line exactly as previously described [18]. Individual gene overexpression and gene targeting experiments were performed as previously reported using previously characterized reagents and cell lines [17; 18].

## Statistical Analysis

For single-cell analysis individual cells were considered as biological replicates (n = 241). Calculated primer efficiencies revealed a normal distribution determined by the Shapiro-Wilk test. For normal distributed data we used the two-tailed Student’s t-test for significance calculations. Nonparametric statistical approaches were applied for data not following a normal distribution. Specifically, we chose the Kurskal-Wallis test for independent and unequal sized sample calculations. Statistical significance was set to p < 0.05 for gene expression analysis (n > 3). Only Bayesian network connections with p < 0.05 are shown. Correlation analysis revealed only significant correlations with p < 0.05. Resulting values (for each experiment) were subject to two-tailed Student’s t-test and two-tailed Fisher’s exact test. Error bars represent SEM in all statistical significance tests. Follow-up validation analyses were performed using five independent validation data sets of single-cell genome-wide expression profiling analyses of human and mouse preimplantation embryogenesis (Supplemental Table S1), including genome-wide expression profiles of 1,708 individual human cells [12; 13; 34; 44] and 259 individual mouse cells [45].

## Author contributions

J.D.D. and M.W. designed, performed, and interpreted experiments. G.V.G. performed the HPAT lincRNAs expression profiling analyses, HPAT lincRNA expression-defined identification of human blastocyst sub-populations, follow-up validation analyses of gene expression profiles of individual human embryonic cells, and formulated a two-wave embryogenesis progression model. G.V.G, V.S. and J.D.D. contributed to the analyses and interpreting results. G.V.G., J.D.D., M.W, and V.S. wrote and edited the manuscript.

## Acknowledgments

We thank Stefan Heller and Joanna Wysocka for thoughtful feedback and comments regarding data analysis and manuscript preparation. We thank Rudolph Jaenisch for providing the OCT4-ΔPE-GFP line. We thank Toshinobu Nishimura for providing the PiggyBac constructs. This work was funded by the grants: U54-1U54HD068158-01 (Stanford University Center for Reproductive and Stem Cell Biology), RB3-2209 (California Institute of Regenerative Medicine), start-up funds to V.S. V.S. is also supported by Siebel Stem Cells Scholarship. J.D.D. is supported by Fritz Thyssen Fellowship.

## References

1. Santoni, F.A., Guerra, J., and Luban, J. HERV-H RNA is abundant in human embryonic stem cells and a precise marker for pluripotency. Retrovirology 2012; 9, 111.

2. Xie W, Schultz MD, Lister R, Hou Z, Rajagopal N, Ray P, Whitaker JW, Tian S, Hawkins RD, Leung D, Yang H, Wang T, Lee AY, Swanson SA, Zhang J, Zhu Y, Kim A, Nery JR, Urich MA, Kuan S, Yen CA, Klugman S, Yu P, Suknuntha K, Propson NE, Chen H, Edsall LE, Wagner U, Li Y, Ye Z, Kulkarni A, Xuan Z, Chung WY, Chi NC, Antosiewicz-Bourget JE, Slukvin I, Stewart R, Zhang MQ, Wang W, Thomson JA, Ecker JR, Ren B. Epigenomic analysis of multilineage differentiation of human embryonic stem cells. Cell 2013. 153, 1134–1148.

3. Glinsky, G.V. Transposable Elements and DNA methylation create in embryonic stem cells human-specific regulatory sequences associated with distal enhancers and noncoding RNAs. Genome Biol Evol. 2015; 7: 1432–54.

4. Kelley, D., and Rinn, J. Transposable elements reveal a stem cell-specific class of long noncoding RNAs. Genome Biol. 2012; 13, R107.

5. Kunarso, G., Chia, N.Y., Jeyakani, J., Hwang, C., Lu, X., Chan, Y.S., Ng, H.H., and Bourque, G. Transposable elements have rewired the core regulatory network of human embryonic stem cells. Nat Genet. 2010; 42, 631–634.

6. Smith ZD, Chan MM, Humm KC, Karnik R, Mekhoubad S, Regev A, Eggan K, Meissner A. DNA methylation dynamics of the human preimplantation embryo. Nature 2014; 511: 611–615.

7. Fort A, Hashimoto K, Yamada D, Salimullah M, Keya CA, Saxena A, Bonetti A, Voineagu I, Bertin N, Kratz A, Noro Y, Wong CH, de Hoon M, Andersson R, Sandelin A, Suzuki H, Wei CL, Koseki H; FANTOM Consortium, Hasegawa Y, Forrest AR, Carninci P. Deep transcriptome profiling of mammalian stem cells supports a regulatory role for retrotransposons in pluripotency maintenance. Nature Genet. 2–14; 46: 558–566.

8. Lu X, Sachs F, Ramsay L, Jacques PÉ, Göke J, Bourque G, Ng HH. The retrovirus HERVH is a long noncoding RNA required for human embryonic stem cell identity. Nat Struct Mol Biol. 2014; 21:423–425.

9. Ohnuki M, Tanabe K, Sutou K, Teramoto I, Sawamura Y, Narita M, Nakamura M, Tokunaga Y, Nakamura M, Watanabe A, Yamanaka S, Takahashi K. Dynamic regulation of human endogenous retroviruses mediates factor-induced reprogramming and differentiation potential. Proc Natl Acad Sci USA. 2014. 111:12426–31.

10. Koyanagi-Aoi M, Ohnuki M, Takahashi K, Okita K, Noma H, Sawamura Y, Teramoto I, Narita M, Sato Y, Ichisaka T, Amano N, Watanabe A, Morizane A, Yamada Y, Sato T, Takahashi J, Yamanaka S. Differentiation-defective phenotypes revealed by large-scale analyses of human pluripotent stem cells. Proc Natl Acad Sci USA. 2013; 110: 20569–74.

11. Marchetto MC, Narvaiza I, Denli AM, Benner C, Lazzarini TA, Nathanson JL, Paquola AC, Desai KN, Herai RH, Weitzman MD, Yeo GW, Muotri AR, Gage FH. (2013). Differential LINE-1 regulation in pluripotent stem cells of humans and other great apes. Nature 503: 525–529.

12. Xue Z, Huang K, Cai C, Cai L, Jiang CY, Feng Y, Liu Z, Zeng Q, Cheng L, Sun YE, Liu JY, Horvath S, Fan G. Genetic programs in human and mouse early embryos revealed by single-cell RNA sequencing. Nature 2013; 500: 593–597.

13. Yan L, Yang M, Guo H, Yang L, Wu J, Li R, Liu P, Lian Y, Zheng X, Yan J, Huang J, Li M, Wu X, Wen L, Lao K, Li R, Qiao J, Tang F. Single-cell RNA-Seq profiling of human preimplantation embryos and embryonic stem cells. Nat Struct Mol Biol 2013; 20: 1131–1139.

14. Goke J, Lu X, Chan YS, Ng HH, Ly LH, Sachs F, Szczerbinska I. Dynamic transcription of distinct classes of endogenous retroviral elements marks specific populations of early human embryonic cells. Cell Stem Cell 2015; 16: 135–141.

15. Wang J, Xie G, Singh M, Ghanbarian AT, Raskó T, Szvetnik A, Cai H, Besser D, Prigione A, Fuchs NV, Schumann GG, Chen W, Lorincz MC, Ivics Z, Hurst LD, Izsvák Z. Primate-specific endogenous retrovirus-driven transcription defines naïve-like stem cells. Nature 2014; 516: 405–9.

16. Grow EJ, Flynn RA, Chavez SL, Bayless NL, Wossidlo M, Wesche DJ, Martin L, Ware CB, Blish CA, Chang HY, Pera RA, Wysocka J. Intrinsic retroviral reactivation in human preimplantation embryos and pluripotent cells. Nature 2015; 522: 221–5.

17. Durruthy-Durruthy J, Sebastiano V, Wossidlo M, Cepeda D, Cui J, Grow EJ, Davila J, Mall M, Wong WH, Wysocka J, Au KF, Reijo Pera RA. The primate-specific noncoding RNA HPAT5 regulates pluripotency during human preimplantation development and nuclear reprogramming. Nat Genet. 2016; 48: 44–52.

18. Durruthy-Durruthy J, Wossidlo M, et al. Spatiotemporal reconstruction of the human blastocyst by single-cell gene expression analysis informs induction of naive pluripotency. Dev Cells 2016; 38: 100–115.

19. Vanneste E, Voet T, Le Caignec C, Ampe M, Konings P, Melotte C, Debrock S, Amyere M, Vikkula M, Schuit F, Fryns JP, Verbeke G, D’Hooghe T, Moreau Y, Vermeesch JR. Chromosome instability is common in human cleavage-stage embryos. Nat Med. 2009; 15: 577–83.

20. Johnson DS, Gemelos G, Baner J, Ryan A, Cinnioglu C, Banjevic M, Ross R, Alper M, Barrett B, Frederick J, Potter D, Behr B, Rabinowitz M. Preclinical validation of a microarray method for full molecular karyotyping of blastomeres in a 24-h protocol. Hum Reprod. 2010; 25: 1066–75.

21. Chavez SL, Loewke KE, Han J, Moussavi F, Colls P, Munne S, Behr B, Reijo Pera RA. Dynamic blastomere behaviour reflects human embryo ploidy by the four-cell stage. Nat Commun. 2012; 3:1251.

22. Vera-Rodriguez M, Chavez SL, Rubio C, Reijo Pera RA, Simon C. Prediction model for aneuploidy in early human embryo development revealed by single-cell analysis. Nat Commun. 2015; 6: 7601.

23. Yanez LZ, Han J, Behr BB, Pera RA, Camarillo DB. Human oocyte developmental potential is predicted by mechanical properties within hours after fertilization. Nat Commun. 2016; 7: 10809.

24. Arman, E., Haffner-Krausz, R., Chen, Y., Heath, J.K., and Lonai, P. (1998). Targeted disruption of fibroblast growth factor (FGF) receptor 2 suggests a role for FGF signaling in pregastrulation mammalian development. Proceedings of the National Academy of Sciences of the United States of America 95, 5082–5087.

25. Brevini, T.A.L., and Pennarossa, G. (2013). Gametogenesis, early embryo development and stem cell derivation (New York, NY: Springer).

26. Nichols, J., Silva, J., Roode, M., and Smith, A. (2009). Suppression of Erk signalling promotes ground state pluripotency in the mouse embryo. Development 136, 3215–3222.

27. Feldman, B., Poueymirou, W., Papaioannou, V.E., DeChiara, T.M., and Goldfarb, M. (1995). Requirement of FGF-4 for postimplantation mouse development. Science 267, 246–249.

28. Chazaud, C., Yamanaka, Y., Pawson, T., and Rossant, J. (2006). Early lineage segregation between epiblast and primitive endoderm in mouse blastocysts through the Grb2-MAPK pathway. Developmental Cell 10, 615–624.

29. Cheng, A.M., Saxton, T.M., Sakai, R., Kulkarni, S., Mbamalu, G., Vogel, W., Tortorice, C.G., Cardiff, R.D., Cross, J.C., Muller, W.J., et al. (1998). Mammalian Grb2 regulates multiple steps in embryonic development and malignant transformation. Cell 95, 793–803.

30. Takashima, Y., Guo, G., Loos, R., Nichols, J., Ficz, G., Krueger, F., Oxley, D., Santos, F., Clarke, J., Mansfield, W., et al. (2014). Resetting transcription factor control circuitry toward ground-state pluripotency in human. Cell 158, 1254–1269.

31. Theunissen, T.W., Powell, B.E., Wang, H., Mitalipova, M., Faddah, D.A., Reddy, J., Fan, Z.P., Maetzel, D., Ganz, K., Shi, L., et al. (2014). Systematic identification of culture conditions for induction and maintenance of naive human pluripotency. Cell Stem Cell 15, 471–487.

32. Qin H, Hejna M, Liu Y, Percharde M, Wossidlo M, Blouin L, Durruthy-Durruthy J, Wong P, Qi Z, Yu J, Qi LS, Sebastiano V, Song JS, Ramalho-Santos M. (2016). YAP induces human naive pluripotency. Cell Rep. 14: 2301–12.

33. Lin DY, Shih HM. (2002). Essential role of the 58-kDa microspherule protein in the modulation of Daxx-dependent transcriptional repression as revealed by nucleolar sequestration. J Biol Chem. 277: 25446–56.

34. Petropoulos S, Edsgärd D, Reinius B, Deng Q, Panula SP, Codeluppi S, Plaza Reyes A, Linnarsson S, Sandberg R, Lanner F. (2016). Single-Cell RNA-Seq Reveals Lineage and X Chromosome Dynamics in Human Preimplantation Embryos. Cell 165: 1012–26.

35. Glinsky GV. (2016). Activation of endogenous human stem cell-associated retroviruses (SCARs) and therapy-resistant phenotypes of malignant tumors. Cancer Lett. 376: 347–59.

36. Au, K.F. et al. Characterization of the human ESC transcriptome by hybrid sequencing. (2013). Proc. Natl. Acad. Sci USA 110, E4821–E4830.

37. Huang, K., Maruyama, T., and Fan, G. (2014). The naive state of human pluripotent stem cells: a synthesis of stem cell and preimplantation embryo transcriptome analyses. Cell Stem Cell 15: 410–415.

38. Treff NR, Su J, Taylor D, Scott Jr. RT. (2011). Telomere DNA deficiency is associated with development of human embryonic aneuploidy. PLoS genetics. 7: e1002161.

39. Ulaner GA, Giudice LC. (1997). Developmental regulation of telomerase activity in human fetal tissues during gestation. Mol Hum Reprod. 3: 769–73.

40. Wright WE, Piatyszek MA, Rainey WE, Byrd W, Shay JW. (1996). Telomerase activity in human germline and embryonic tissues and cells. Dev Genet. 18: 173–9.

41. Cockburn, K., and Rossant, J. (2010). Making the blastocyst: lessons from the mouse. J. Clin. Invest. 120, 995–1003.

42. Yamanaka, Y., Lanner, F., and Rossant, J. (2010). FGF signal-dependent segregation of primitive endoderm and epiblast in the mouse blastocyst. Development 137, 715–724.

43. Nikas, G., Ao, A., Winston, R.M., and Handyside, A.H. (1996). Compaction and surface polarity in the human embryo in vitro. Biol. Reprod. 55, 32–37.

44. Dang, Y. et al. (2016). Tracing the expression of circular RNAs in human pre-implantation embryos. Genome Biol. 17: 130.

45. Deng, Q., Ramskold, D., Reinius, B., and Sandberg, R. (2014). Single-cell RNA-seq reveals dynamic, random monoallelic gene expression in mammalian cells. Science 343: 193–196.

46. Wong CC, Loewke KE, Bossert NL, Behr B, De Jonge CJ, Baer TM, Reijo Pera RA. (2010). Non-invasive imaging of human embryos before embryonic genome activation predicts development to the blastocyst stage. Nat Biotechnol. 28: 1115–21.

47. Holt SE, Glinsky VV, Ivanova AB, Glinsky GV. (1999). Resistance to apoptosis in human cells conferred by telomerase function and telomere stability. Mol Carcinog. 25: 241–8.

48. Glinsky GV. (2016). Single cell genomics reveals activation signatures of endogenous SCAR’s networks in aneuploid human embryos and clinically intractable malignant tumors. Cancer Lett. 381: 176–193. doi: 10.1016/j.canlet.2016.08.001. PMID: 27497790.

49. Das A, Ghosh J, Bhattacharya A, Samanta D, Das D, Das Gupta C. (2011). Involvement of mitochondrial ribosomal proteins in ribosomal RNA-mediated protein folding. J Biol Chem. 286: 43771–81.

